# Competition for space induces cell elimination through compaction-driven ERK downregulation

**DOI:** 10.1101/447888

**Authors:** Eduardo Moreno, Léo Valon, Florence Levillayer, Romain Levayer

**Affiliations:** Champalimaud Centre for the Unknown, Av. Brasília 1400-038, Lisbon, Portugal; Institut Pasteur, Department of Developmental and Stem Cell Biology, 25 rue du Dr. Roux, 75015 Paris, France

**Keywords:** mechanotransduction, cell competition, EGFR, ERK, Hid, epithelium

## Abstract

The plasticity of developing tissues relies on the adjustment of cell survival and growth rate to environmental cues. This includes the effect of mechanical cues on cell survival. Accordingly, compaction of an epithelium can lead to cell extrusion and cell death. This process was proposed to contribute to tissue homeostasis but also to facilitate the expansion of pretumoral cells through the compaction and elimination of the neighbouring healthy cells. However we know very little about the pathways than can trigger apoptosis upon tissue deformation and the contribution of compaction driven death to clone expansion was never assessed *in vivo*. Using the *Drosophila* pupal notum and a new live sensor of ERK, we show that tissue compaction induces cell elimination through the downregulation of EGFR/ERK pathway and the upregulation of the pro-apoptotic protein Hid. Those results suggest that the sensitivity of EGFR/ERK pathway to mechanics could play a more general role in the fine tuning of cell elimination during morphogenesis and tissue homeostasis. Secondly, we assessed *in vivo* the contribution of compaction driven death to pretumoral cell expansion. We found that the activation of the oncogene Ras in clones can also downregulate ERK and activate apoptosis in the neighbouring cells through their compaction, which contributes to Ras clone expansion. The mechanical modulation of EGFR/ERK during growth-mediated competition for space may contribute to tumour progression.

## Introduction

Developing tissues have the capacity to cope with perturbations, which include stress from the environment (mechanical, chemical, temperature) as well as abnormal behaviors of a subset of cells (misspecification, local modification of growth rate). This robustness relies on the plasticity of cell behavior/fate which can be adjusted to changes in the tissue environment. The adjustment of cell proliferation and cell death could rely on several inputs, including contact dependent communication, communication through extracellular diffusive factors and/or communication through mechanical inputs[1]. Accordingly, mechanical cues have been proposed to adjust the local rate of cell death and cell division to regulate the final size of a tissue or to maintain constant size during homeostasis [2-4].

The regulation of cell death by mechanical inputs is well documented in epithelia. Compaction of epithelia was shown to induce cell extrusion and cell death in MDCK cells, zebrafish [5-7] and in the midline region of the *Drosophila* pupal notum (a single layer epithelium) [8] (**Fig. 1A**). Recently, we showed that compaction driven cell elimination in the pupal notum relies on caspase activation which was required for and preceding every extrusion event [9]. This suggested that some unknown pathways could be sensitive to tissue deformations and trigger caspase activation. However, we could not find a clear contribution of known mechanosensitive pathways to midline cell elimination, including the JNK pathway[10] or the Hippo Yap/Taz pathway [9, 11]. Therefore, the pathway regulating cell elimination in the midline still remained unknown.

**Figure 1:**
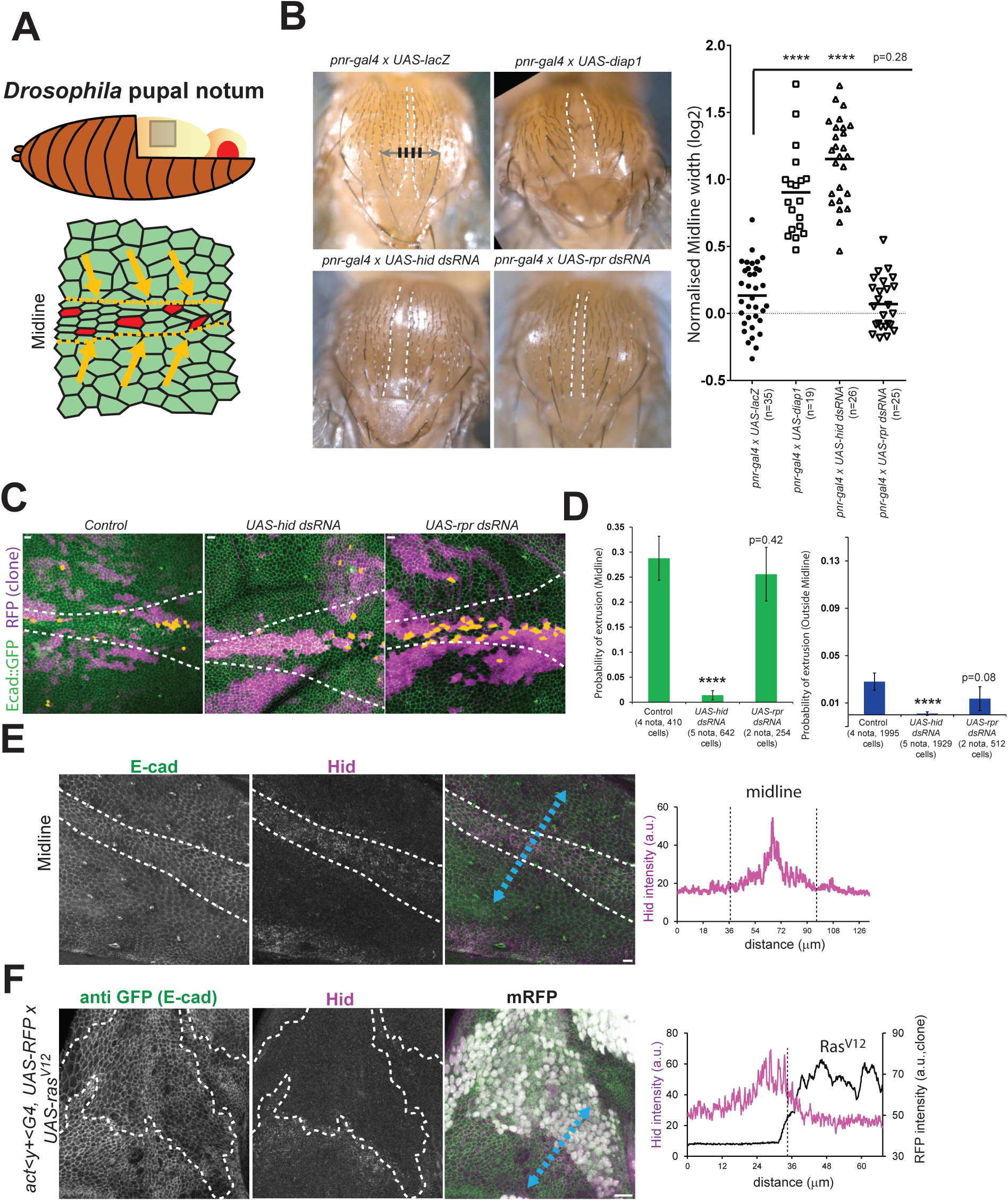
Hid is required for cell elimination in the pupal notum (associated with movie S1) **A:** Schematic of the pupal notum. Bottom shows the midline region where cells undergo compaction (orange arrow) and where the rate of cell elimination is high (red cells=caspase activated cells). **B:** Representative adult thorax upon perturbation of cell death in the *pnr* domain. White dashed lines encompassed the midline. Grey line is the line connecting the two aDC macrochaetae used to measure the width of the midline and the two surrounding bristle rows (black lines). Right graph is the normalised midline width (log2 scale) in every condition (one point=1 thorax), p-values are t-test with the control, ****=p<10^-4^. **C:** z-projection of live pupal nota expressing *ubi-Ecad::GFP* (green) with Gal4 expressing clones (RFP, magenta) in controls (ayG4 alone) or expressing *UAS-hid dsRNA* or *UAS-rpr dsRNA*. The midline is surrounded by white dashed lines. Orange cells are the clonal cells that will die over the course of the movie (700 min). Scale bars=10μm. **D:** Quantification of the probability of cell elimination in the clones in the midline region (left) and outside the midline region (right). p-values are Fisher exact test with the control condition. ****=p<10^-4^. Note that we did the same experiment with *UAS-grim dsRNA* (GD21830) and observed a significant increase of the rate of cell death in and outside the midline. However all the UAS-grim dsRNA line available have multiple off targets (not shown). **E:** Immunostaining of a pupal notum showing z-projection of anti E-cad (green) and anti Hid (magenta) in the midline region (white dashed line) (7/7 nota). Right graph shows the intensity profile of Hid along the blue dashed line (magenta). Scale bar=10μm. **F:** Immunostaining of a pupal notum showing z-projection of anti GFP (E-cad::GFP, green), anti Hid (magenta) and UAS-nlsRFP signal (white) in vicinity of a clone where Ras was conditionally activated (*UAS-ras^V12^*, 24h activation at 29°C) (white dashed lines show clone boundaries), 4/4 nota. Right graph shows the intensity profile of Hid along the blue dashed line (magenta) and the RFP signal marking the clone (black line). Note that Hid accumulated up to 10-15μm (2-3 cells) away from the clone. Scale bar=10μm.

The contribution of caspases to cell elimination also suggested that cells could have a differential sensitivity to compaction depending on their sensitivity to apoptosis. Accordingly, activation of Ras in clones led to the preferential compaction and elimination of the neighbouring wild type (WT) cells [9]. Similarly, the high levels of p53 in mutant MDCK cells for the polarity gene *scribble* increase their sensitivity to compaction and trigger their preferential elimination when surrounded by WT MDCK cells [7]. Therefore those mechanical driven deaths have been proposed to promote the expansion of pretumoural cells through a so called mechanical cell competition that drives preferential elimination of the neighbouring healthy cells [7, 9, 12, 13]. However, the molecular pathway triggering cell death during mechanical cell competition *in vivo* was not yet identified and it was not yet clear whether such elimination could significantly promote pretumoural clone expansion.

Here, we show that tissue compaction induces cell elimination in the pupal notum through the downregulation of Epidermal Growth Factor Receptor/Extracellular signal Regulated Kinase (EGFR/ERK) pathway and the upregulation of the pro-apoptotic protein Hid (Head Involution Defective). Using a new *Drosophila* live sensor of ERK activity, we demonstrate that local tissue stretching/compaction can transiently upregulate/downregulate ERK activity hence increasing/decreasing cell survival. Moreover, we show that compaction driven ERK downregulation near Ras activated clones controls cell elimination and promotes clone expansion. The sensitivity of EGFR/ERK pathway to mechanics and its role in the fine tuning of cell elimination could play a more general role during tissue homeostasis and tumour progression

## Results

### Cell elimination in the pupal notum is regulated by Hid

We previously showed that a deletion covering the three pro-apoptotic genes *hid*, *grim* and *reaper* (*H99* deletion) strongly downregulated cell extrusion in the pupal notum [9]. We therefore tried to identify which of those genes was required for cell elimination. Downregulation of *hid* by RNAi in the pupal notum (using *pnr-gal4* driver) led to a significant widening of the midline region in the adult fly thorax (the midline is a zone with a high rate of cell elimination [8, 9, 14]), similar to apoptosis downregulation through *diap1* overexpression (**Fig1.B**). Accordingly, downregulation of *hid* in clones strongly downregulated their rate of cell extrusion whether inside or outside the midline (**Fig1.C,D, movie S1**), phenocopying the defects observed with the *H99* deletion[9]. This suggested that *hid* was responsible for the defects observed in *H99* deletion. Accordingly, *rpr* downregulation had no significant effect on thorax morphology and clone survival (**Fig 1B-D movie S1**). Finally, we observed an accumulation of Hid protein in regions showing a high rate of cell elimination, including in the midline region (**Fig. 1E**) and in crowded regions near *Ras^V12^* clones (**Fig. 1F**). Altogether, this suggested that Hid is a central regulator of cell death in the pupal notum and that it could be upregulated in crowded regions.

### Cell survival in the midline is regulated by the EGFR/Ras/Raf/ERK pathway

We next tried to identify the potential regulators of Hid in the midline region of the notum. Several regulators of Hid were previously described in *Drosophila*, including upregulation by the steroid hormone Ecdysone [15], downregulation by the pro-survival microRNA Bantam [16], and downregulation by the EGFR/Ras/Raf/ERK pathway [17, 18]. We did not observe any accumulation of Ecdysone receptor (**Fig. S1A**) nor a downregulation of Bantam in the midline region (**Fig. S1B**, a slight reduction of the GFP signal in the midline rather suggested an increase of Bantam activity), suggesting that they were not responsible for the upregulation of Hid in the midline. Moreover, overexpression of a Bantam sponge in clones (which sequesters Bantam microRNA[16], and can significantly reduce Bantam activity, **Fig. S1C**) had no significant effect on their rate of extrusion (**Fig. S1D,E, movie S2**). However, we observed a downregulation of ERK in the midline region using diphosphoERK staining (dpERK, **Fig. S1F**) and the nuclear accumulation of Capicua (negatively regulated by ERK[19]) (**Fig. 2A**). As such, we tried to identify the upstream signal regulating ERK activity in the midline. Downregulation of EGFR in the thorax using *pnr-gal4* driver led to a narrowing of the *pnr* domain in the adult (domain encompassed by the two large macrochaetes, central white doted line, **Fig. 2B**). This effect was driven by the upregulation of *hid* dependent death as the phenotype could be rescued by downregulation of *hid* (**Fig. 2B**). Moreover, it was specific of EGFR as we did not observe such narrowing upon Pvr/PDGF, FAK, Src42 and Src64 downregulation (**Fig. S2A**), other putative regulators of ERK[20-22]. We then tested the impact of EGFR on cell survival in the notum. Downregulation of EGFR in clones led to a strong increase of the cell extrusion rate both in the midline and outside the midline while overexpression of a chimeric EGFR (which can activate ERK independently of extracellular ligands[23]) nearly abolished cell extrusion (**Fig. 2C,D, movie S3,** the effect of UAS-hEGFR::GFP and EGFRdsRNA on ERK is shown in **Fig. 3C**, see below). This was suggesting that EGFR pathway downregulation was necessary and sufficient for cell elimination in the notum. EGFR/Ras could promote cell survival through the PI3K pathway or the Raf/MAPK pathway[24]. However activation of PI3K in clones had little impact on the rate of cell elimination (**Fig. S2B, movie S2**) while overexpression of an active form of Raf strongly abolished cell elimination (**Fig. S2C, movie S2**). Altogether, this was suggesting that EGFR/Ras/Raf/ERK pathway was a central regulator of cell survival in the pupal notum.

**Figure 2:**
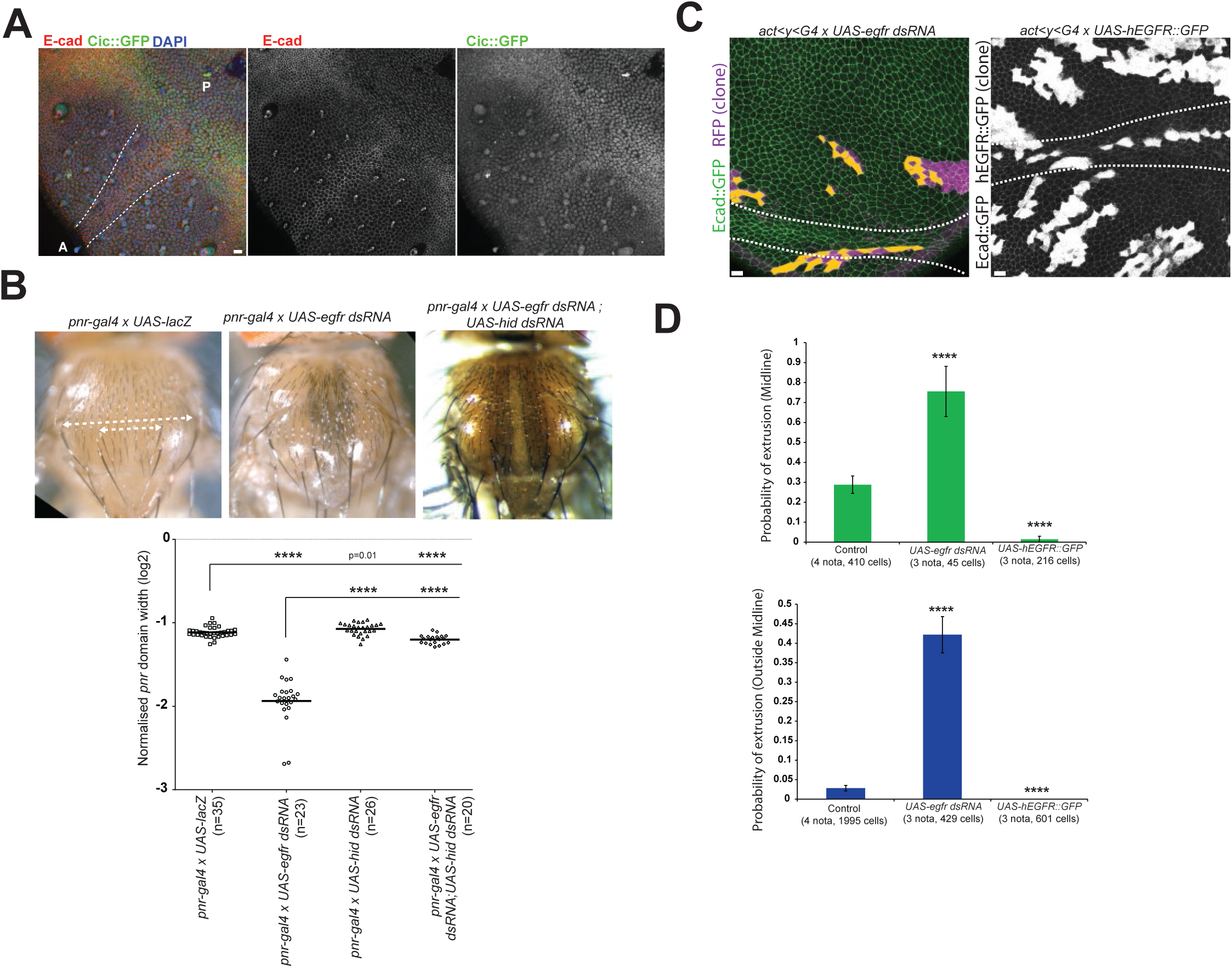
EGFR/ERK downregulation is necessary and sufficient for cell elimination (associated with movie S3) **A:** z-projection of an immunostaining of a pupal notum, anti E-cad (red), DAPI (blue) and anti GFP (*cic-Cic::GFP*, BAC clone, green) in the midline region (white dashed line) (7/7 nota). A: Anterior, P: Posterior. 5/5 nota. Scale bar=10μm. **B:** Representative adult thorax in control (*pnr-gal4 x UAS-lacZ)* upon EGFR downregulation (*pnr-gal4* x *UAS-EGFR dsRNA*) and downregulation of EGFR and Hid (*pnr-gal4* x *UAS-EGFR dsRNA; UAS-hid dsRNA*). White dashed lines show the *pnr* domain width (distance between two aDC macrochaetae) and total thorax width (distance between two pSA macrochaetae). Note that Hid downregulation rescued the reduction of size of the *pnr* domain, but not the pigmentation defects. Bottom graph is the normalised *pnr* domain width (log2 scale) in every condition (one point=1 thorax), pvalues are t-tests with the control or *UAS-egfr dsRNA*, ****=p<10^-4^. **C:** z-projection of live pupal nota expressing *ubi-Ecad::GFP* (green) with Gal4 expressing clones where EGFR is downregulated (*UAS-egfr dsRNA*, left, clones in magenta), or upregulated (*UAS-hEGFR::GFP*, *h. sapiens* extracellular domain and *Drosophila* intracellular domain, clones marked in white). The midline is delineated by white dashed lines. Orange cells are the clonal cells that will die over the course of the movie (700 min). See **Fig. 1C** for control. Scale bars=10μm. **D:** Quantification of the probability of cell elimination in the clones in the midline region (top) and outside the midline region (bottom). p-values are Fisher exact test with the control condition (same as **Fig. 1C,D**). ****=p<10^-4^.

### Developmental of a new live sensor of ERK

We then asked whether ERK downregulation was playing an instructive role for caspase activation rather than a purely permissive function. To do so, we developed a new live sensor of ERK activity. Phosphorylation of the transcriptional repressor Capicua (Cic) by ERK leads to its removal from the nucleus[19]. We used a minimal domain of Cic lacking its DNA binding domain and containing the phosphorylation and docking site of ERK[19] fused to a NLS and mCherry under the control of the *tubulin* promoter (**Fig. 3A**, hereafter named miniCic). We expected to observe an accumulation of nuclear miniCic upon ERK downregulation and an exclusion from the nucleus upon ERK activation. miniCic nuclear intensity was indeed anticorrelated with the endogenous levels of active ERK in the wing and eye imaginal discs (**Fig. 3B**) and in the embryos (**Fig. S3**). Note that the apparent low signal of miniCic in active ERK regions is due to the dilution of the signal in the full cytoplasm of the cells. The variations of miniCic intensity along ERK gradient (**Fig. 3B** gradient in the eye disc[25], bottom graph) suggested that miniCic could be used as a quantitative proxy for endogenous levels of ERK activity. Moreover, miniCic could accurately report modulations of EGFR/ERK activity in clones in the pupal notum. Accordingly, reduction of ERK activity in clones through EGFR depletion by RNAi in regions of normally high ERK activity (lateral regions) led to a clear ectopic miniCic nuclear accumulation (**Fig. 3C** left) while activation of ERK in clones through hEGFR::GFP expression in regions of normally low ERK activity (posterior region) led to a clear ectopic exclusion of miniCic from the nucleus (**Fig. 3C** right). Thus, miniCic can be used to measure the levels of ERK activity in the pupal notum. We then assessed the potential time delay between ERK modulation and miniCic relocalisation. Inhibition of ERK through the MEK inhibitor Trametinib[26] led to an almost immediate disappearance of dpERK in S2 cells (**Fig. 3D**, 5min post drug treatment at 10μM). Similarly, treatment of haemocytes expressing miniCic with Trametinib led to a rapid accumulation of miniCic in the nucleus with a significant increase 10 min after drug treatment (**Fig. 3E, movie S4**). Moreover the fast induction of dpERK in the lateral ectoderm of cellularising embryos[27] was also captured by miniCic, except for a short developmental time window of 10-15min (**Fig. S3**, developmental time estimated by the length of the invaginating membrane furrows[27]). Altogether, this showed that miniCic could be used to assess the variations of activity of endogenous ERK with a maximum timelag of 10-15 min. We therefore used this new sensor to visualize the dynamics of ERK in the pupal notum. As suggested by immunostainings (**Fig. 2A, Fig. S1F**), we observed an accumulation of miniCic (low ERK activity) mostly in the midline region and in an orthogonal domain in the posterior part of the notum (**Fig. 3F**, **movie S5**), both corresponding to zones with high rates of cell death[9, 14]. Interestingly, we also observed complex dynamics of ERK activity with global transient downregulation (16-17h after pupal formation, APF) or upregulation (20h APF) in the tissue (**Movie S5**) that respectively followed and preceded the first and second waves of divisions. This suggested that ERK might be modulated by multiple factors during pupal development.

**Figure 3:**
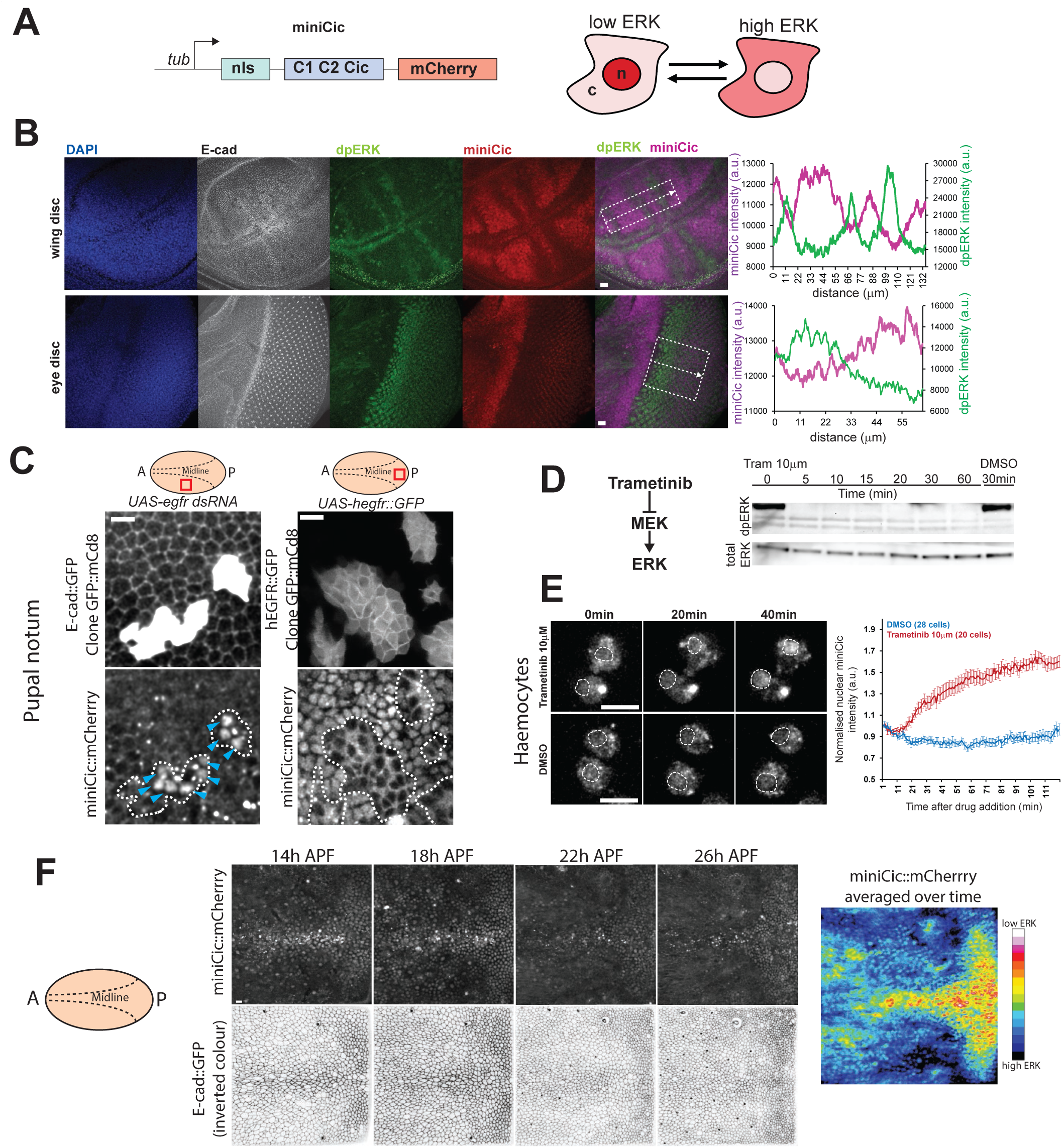
A new live sensor of ERK (associated with Fig. S3 and movie S4 and S5) **A:** Schematic of the miniCic construct. *tubulin* promoter drives the expression of a nls fused to C1 and C2 domains of Cic and mCherry in Cter. The right cartoon shows the expected localisation of miniCic at low ERK activity (accumulation in nucleus, n), and high ERK activity (exclusion in the cytoplasm, c). Red levels represent relative miniCic concentrations. **B:** zprojection of immunostainings of a wing imaginal disc (top) and an eye imaginal disc (bottom) at the L3 wandering stage (representative of 10 tissues for each) stained for DAPI in blue, E‐ cad white, dpERK green and miniCic (anti mCherry) in red (purple on the overlay). Scale bars=10μm. Note that miniCic nuclear signal is anticorrelated with the bands/gradient of dpERK. The right graphs show intensity line profiles of dpERK (green) and miniCic (purple) along the indicated white dashed rectangles. **C:** z-projection in live pupal nota expressing miniCic showing clones (GFP positive) where EGFR was depleted (left, *UAS-egfr dsRNA*) or overexpressed (right, *UAS-hegfr::GFP*, chimeric EGFR). The top schemes show the localisation of the clones in the notum (red rectangles, A: anterior, P, posterior, dashed lines= midline, zone of high ERK activity on the left, and low ERK activity for the right). Blue arrows point to ectopic nuclear accumulation of miniCic. White dashed lines show clone boundaries. Scale bars=10μm. **D:** Western blot of dpERK (top) and total ERK (bottom) in S2 cells upon inhibition of ERK phosphorylation by Trametinib (10μm, a potent inhibitor of MEK, left scheme) at different time after drug treatment (in minutes). Control band is for DMSO treatment (30min after DMSO release). dpERK band already disappeared 5min after the treatment. Note that similar results were obtained at 1μM. **E:** Live imaging of a primary culture of larval haemocytes expressing tub-miniCic upon treatment with 10μm Trametinib (top) or DMSO (bottom). Time (minutes) is the time after drug deposition. Doted circles mark the nuclei (as detected on transmitted light channel). Scale bars=10μm. Right graph shows the mean normalised miniCic nuclear intensity after drug treatment (two independent experiments for each condition). Error bars are s.e.m‥ **F:** Snapshots of a live pupal notum expressing E-cad::GFP and miniCic (local z-projection) at different time after pupal formation (APF). Anterior is on the left and posterior on the right, midline is in the center (see left scheme). Scale bar=10μm. The right heat map is the averaged miniCic signal over the full duration of the movie which mostly accumulates in the midline and in an orthogonal posterior region.

### Reduction of ERK activity precedes caspase activation

Using miniCic, we first checked whether ERK downregulation did precede the activation of caspases in the pupal midline. Using the FRET caspase sensor Scat3[28] combined with miniCic, we systematically detected the onset of caspase activation in single cells (maximum inflection of the FRET signal) and measured nuclear miniCic before and after caspase activation. We observed a significant downregulation of ERK signaling that preceded for up to one hour the onset of caspase activation (**Fig. 4A-C**, 61% of the caspase activating cells showing a unambiguous increase of miniCic, n=165 cells, 2 pupae, **movie S6**). This pattern is specific of caspase activating cells as quantification of miniCic in randomly selected cells with no caspase activation during the same developmental time window rather led to a slight downregulation of miniCic/upregulation of ERK (**Fig. 4C**, grey curve), probably because of the global progressive downregulation of miniCic signal observed in the midline (**Fig. 3F, movie S5**). Those data confirmed at the single cell level the instructive role of ERK downregulation on caspase activation. Yet, it also suggested that other unidentified factors may also trigger caspase activation in the notum.

**Figure 4:**
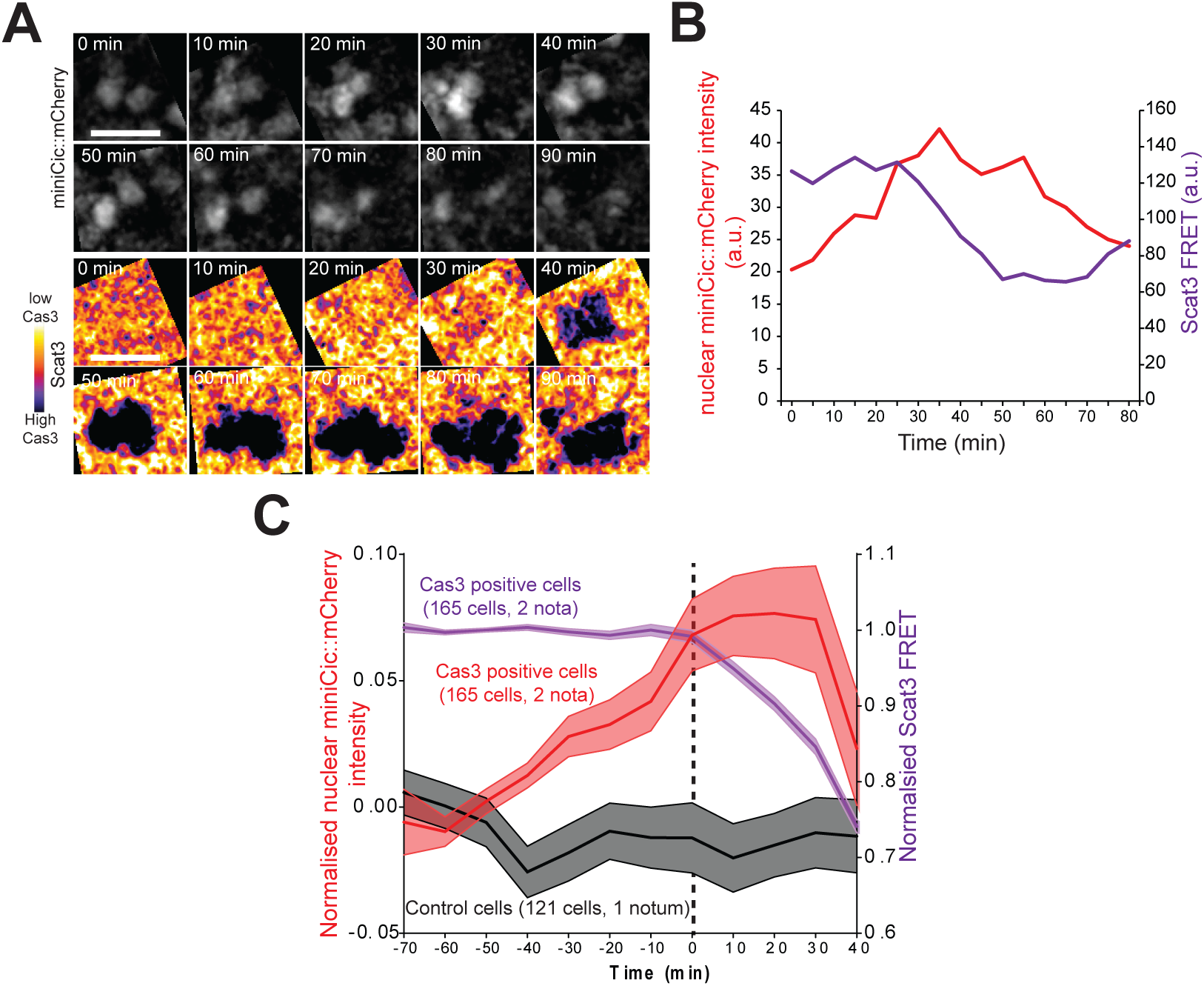
ERK downregulation precedes caspase activation (associated with movie S6) **A:** Snapshots of two nuclei in the midline of the pupal notum showing miniCic signal (top) and Scat3 FRET signal (bottom, dark blue signal= caspase activation), see movie S6. Scale bars=10μm. **B:** miniCic nuclear signal of the left nucleus shown in **A** (red) and the FRET signal (purple). **C:** Averaged normalised profiles of miniCic nuclear intensity (red) in the midline aligned at t0=onset of caspase activation (maximum inflexion of the FRET signal). Light coloured areas are +/- s.e.m‥ The averaged Scat3 FRET signal is shown in purple. The grey curve is the normalised averaged miniCic signal of cells in the same region that do no activate caspase.

### ERK can be ectopically activated/downregulated by tissue stretching/compaction

Cell death in the pupal notum can be downregulated by tissue stretching and ectopically induced by tissue compaction[9]. Since EGFR/ERK is a central regulator of cell survival and cell death in the notum, we wanted to test whether ERK activity could be modulated by tissue deformations. We first tested whether part of the spontaneous dynamics of ERK observed in the notum (**Movie S5**) could be correlated with tissue deformations. We focused on the ERK activation observed around 20h APF (**Movie S5**). Using Particle Image Velocimetry (PIV) to measure local tissue deformations[9], we subdivide the movies in small windows of analysis (11.5x11.5 μm ∼4-5 cells) to correlate the local rate of deformation and local variations of miniCic signal. We found a good correlation between local tissue stretching (**Fig 5A** bottom left panel) and local activation of ERK activity especially in the posterior region of the notum where we observed a global stretch orthogonal to AP axis (**Fig. 5A**,**B movie S7**).This was reflected by the significant positive cross-correlation between miniCic signal derivative and the compaction rate (**Fig 5B,C,** no significant time delay).

**Figure 5:**
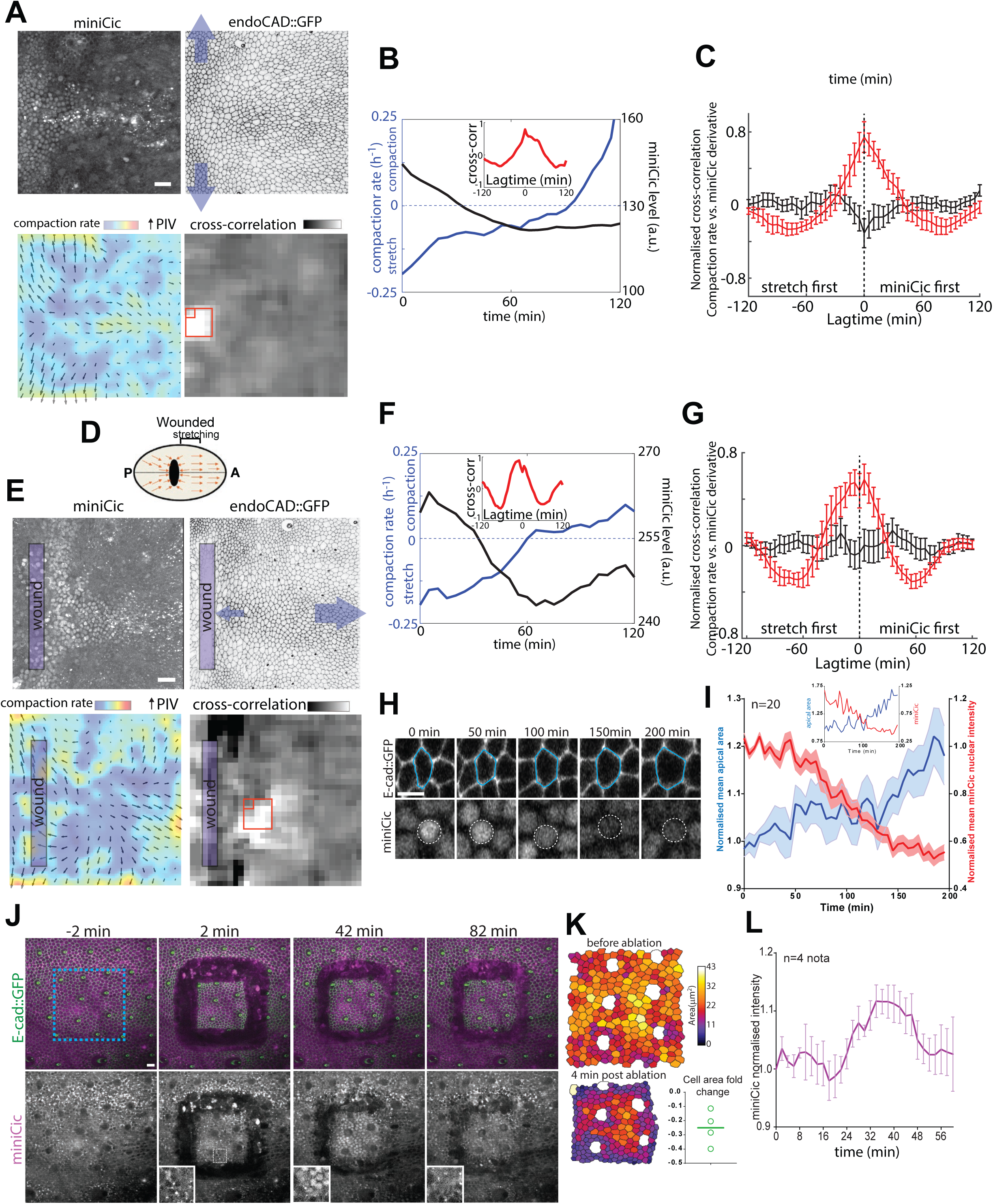
Tissue stretching/compaction can increase/decrease ERK activity ectopically (associated with movies S7,S8,S9) **A:** Imaging of a pupal notum expressing endoCAD::GFP and miniCic::mCherry during a spontaneous phase of tissue stretching orthogonal to the antero-posterior axis (blue arrows, posterior left, anterior right). From left to right and top to bottom, i) miniCic-mCherry fluorescent signal, ii) inverted endoCAD::GFP, iii) overlay of compaction rate and PIV images averaged over 50 minutes (PIV biggest arrow is 5μm/h, compaction rate scale bar ranging from −0.35 to 0.7h^−1^, dark blue=stretching, dark red=compaction). iv) pseudo image of the local cross-correlation value between compaction rate and miniCic signal derivative for a lagtime of 0 minute. Note the peak of correlation that appears in the posterior central region (white region). Scale bar=20μm. **B**: Representative plot of the evolution of the compaction rate (blue) and miniCic levels (black) in a single PIV square (small red rectangle, **Fig. 5A** bottom right panel). Inset, normalised cross-correlation of compaction rate versus the derivative of miniCic for those curves. **C:** Mean normalised cross-correlation of compaction rate versus the derivative of miniCic for the zone of stretching (in red) and control areas (in black) for several nota (n=3) and several regions, errorbars are s.e.m. **D:** Schematic of the wound experiment. The laser wounding (black) generates a zone of stretching anterior to the wound because of the tissue flow to the anterior side (orange arrows) and the wound healing process pulling toward the posterior side. **E:** Imaging of a pupal notum expressing endoCAD::GFP and miniCic::mCherry after laser induced tissue wounding (blue rectangle) inducing ectopic tissue stretching parallel to the antero-posterior axis (blue arrows) anterior to the wound (posterior left, anterior right). From left to right and top to bottom, i) miniCic-mCherry fluorescent signal, ii) inverted endoCAD::GFP, iii) overlay of compaction rate and PIV images averaged over 50 minutes (PIV biggest arrow is 10μm/h, compaction rate scale bar ranging from −0.35 to 0.7h^−1^, dark blue=stretching, dark red=compaction). iv) pseudo image of the local cross-correlation value between compaction rate and miniCic signal derivative for a lagtime of 0 minute. Note the peak of correlation anterior to the wound (white region). Scale bar=20μm. **F**: Representative plot of compaction rate (blue) and miniCic levels (black) in a single PIV square (small red rectangle, **Fig. 5D** bottom right panel). Inset, normalised cross-correlation of compaction rate versus the derivative of miniCic for those curves. **F:** Mean normalised cross-correlation of compaction rate versus the derivative of miniCic for area of stretching (in red) and control areas (in black) for several regions and several nota (n=3), errobars are s.e.m. **H:** Snapshots of a cell in the stretched region anterior to the wound. Cell contour is shown in blue and the corresponding nucleus with miniCic signal with white dotted circles. Scale bar=10μm. **I:** Averaged and normalised evolution of cell apical area and miniCic nuclear signal in the stretched region (n=20 cells). Light colours show s.e.m‥ The inset is the apical area and miniCic nuclear signal of the cell shown in **H**. **J:** Local projection of a live pupal notum expressing endoCAD::GFP (green) and miniCic::mCherry (magenta, bottom) at different times after laser sectioning of a square of 110x10μm (blue square). Insets show a zoom on some nuclei inside the isolated piece of tissue. Scale bar=10μm. **K:** Segmentation of the isolated tissue square prior and post sectioning (apical area in pseudo colours). Bottom right graph shows the average variation of cell apical area (one dot/notum, (A final – A initial)/A initial). **L:** Averaged miniCic intensity in the isolated tissue square (t0=sectioning). Error bars are s.e.m‥

The correlation described above could equally be explained by an effect of ERK on tissue mechanics (as previously reported[29-31]) or an effect of tissue deformation/density on ERK[30, 32-35]. To distinguish between those two possibilities, we tried to ectopically deform the tissue using laser wounding. We previously showed that local laser wounding of the tissue induced cell stretching and reduction of cell elimination anterior to the wound because of the movements driven by the global tissue drift toward the anterior side and the opposite pulling force of the wound closure (see [9], **Fig. 5D**). Accordingly, ectopic stretching of the tissue induced by laser wounding was sufficient to transiently upregulate ERK signaling anterior to the wound (**Fig. 5E,** main stretch along AP axis, **movie S8**). Using PIV, we also observed a significant correlation between zones of stretching and upregulation of ERK activity with no significant time delay (**Fig. 5F,G** positive correlation between compaction rate and miniCic derivative) especially anterior to the wound where most of the stretching occurs (bottom right panel **Fig. 5E**, heat map of the local correlation values). To check whether the local stretching assessed by PIV and the local variations of miniCic intensity did reflect single cell behavior, we tracked cells in the stretched zone, and also observed a significant increase of cell apical area concomitant with a decrease of miniCic nuclear intensity of the corresponding cell (**Fig. 5H,I**). Because the upregulation of ERK and the correlation were restricted to the anterior side of the wound where stretching is stronger (**Fig. 5D,E,** bottom right panel, **movie S8**), this excluded a purely diffusive effect which would act homogenously all around the wound irrespective of the force balance. Altogether, we concluded that tissue/cell stretching could ectopically upregulate ERK activity.

Conversely, we then tested whether ectopic tissue compaction/relaxation could downregulate ERK activity. The pupal notum is globally stretched and laser severing of a piece of epithelium leads to a rapid relaxation of the isolated piece of tissue and reduction the apical area of the cells [36]. Using a pulsed UV laser, we isolated a 110x110μm tissue square (∼300 cells) which rapidly relaxed (**Fig. 5J,K**, **movie S9**). Tissue relaxation/densification was followed by a transient nuclear accumulation of miniCic peaking ∼30min after sectioning (**Fig. 5J,L**, **movie S9**). This suggested that stress release and/or tissue densification could transiently downregulate ERK activity. Interestingly, the effect of wounding on ERK activity in neighbouring cells was opposite in those two laser perturbation experiments (compare **Fig. 5E-I,** activation of ERK and **Fig. 5J-L,** inhibition of ERK), suggesting once again that mechanical inputs rather than diffusive factors from the laser induced wounds were responsible for ERK variations. Altogether, this was showing that tissue stretching/compaction could transiently upregulate/downregulate ERK activity in the pupal notum.

### Compaction driven ERK downregulation is required for cell elimination near Ras^V12^ clones and accelerates clone expansion

We next tested whether ERK activity was also modulated in compacted zones near *Ras^V12^* clones. Induction of ectopic tissue compaction through growth upregulation in *Ras^V12^* clones led to a progressive downregulation of ERK signaling in the neighbouring WT cells (**Fig. 6A**, **movie S10**, region where we normally observed an upregulation of ERK activity, see **Fig 3F**). The pattern of ERK inhibition correlated with the zones of high compaction rate characterised by PIV (**Fig. 6B**, R^2^=0.41) suggesting that ERK activity was specifically downregulated in compacted cells near *Ras^V12^* clones. Moreover, we observed a clear enrichment of apoptotic cells in zones of low ERK activity near Ras clones (**Fig. S4A**). The pattern of ERK downregulation was unlikely to be explained by purely diffusive factors as we did not observe a spatial propagation of ERK downregulation from the clone boundary (**movie S10**), and ERK downregulation was not systematically present near clone boundaries (see for instance **Fig 6A,B**, top left corner, low compaction rate and low miniCic levels).

**Figure 6:**
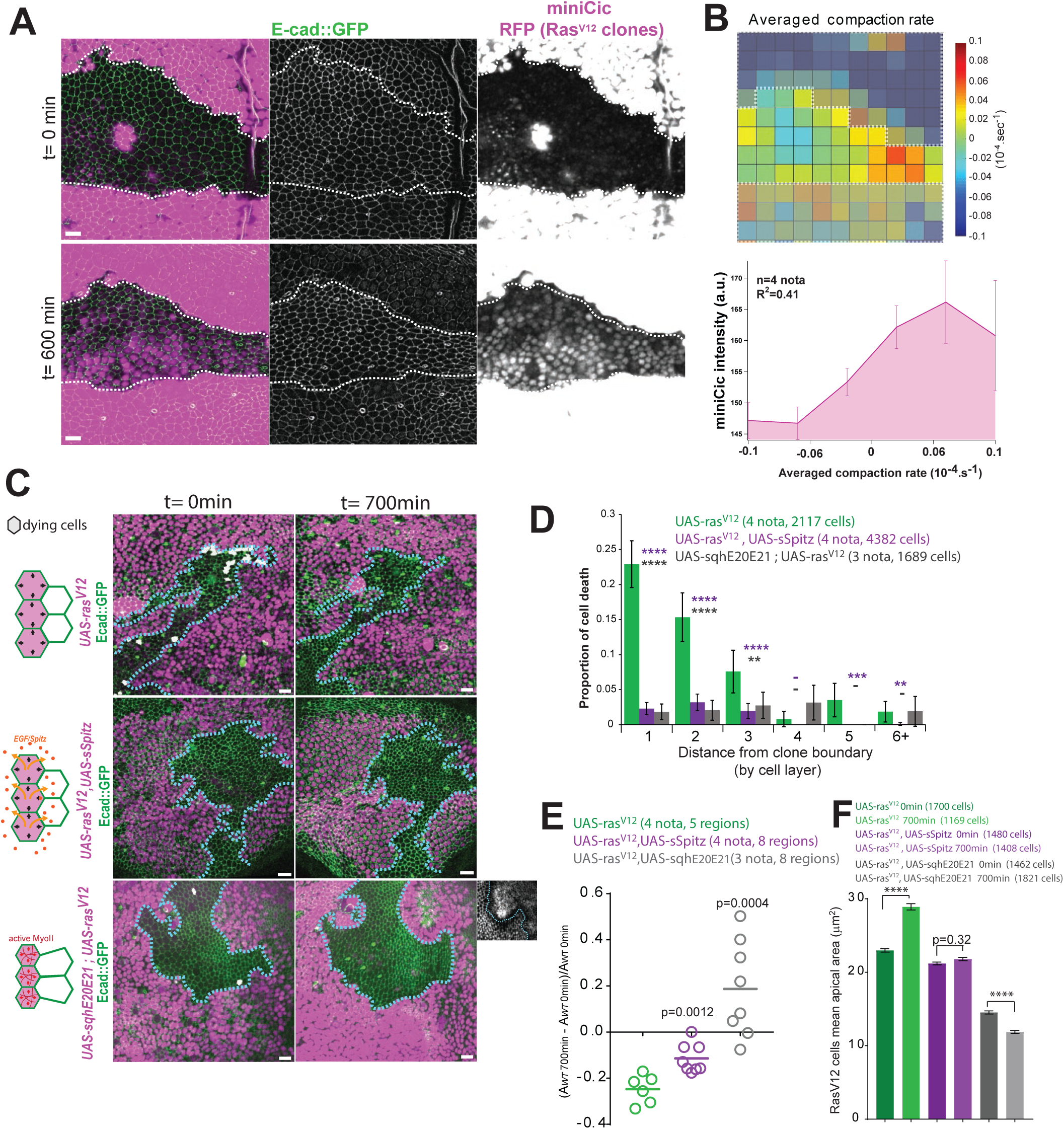
Mechanical competition eliminates cells through compaction driven ERK downregulation (associated with Fig. S4 and movies S10,S11) **A:** Snapshots (0 and 600min, local z-projections) of a live pupal notum expressing endo-Ecad::GFP (green) and miniCic (magenta) upon conditional induction of *Ras^V12^* in clones (RFP, strong magenta signal, white dashed lines). Scale bar=10μm. **B:** Top, Map of the averaged compaction rate calculated by PIV (white dashed lines= clone boundaries). Red=Compaction, Blue=stretching. Bottom, averaged local miniCic intensity at 600min for a given averaged compaction rate (4 nota). R^2^ is the Pearson correlation coefficient. **C:** z-projection (at the onset of the movie and after 700min) of live pupal nota expressing endo-Ecad::GFP (green) (maximum z-projection) upon conditional induction of *Ras^V12^* (left) or *Ras^V12^* and *sSpitz* (*Drosophila* EGF) or *Ras^V12^* and *sqhE20E21* (constitutively active MyosinII Regulatory Light Chain) in clones (RFP, strong magenta signal, blue dashed lines show the contours). The cells that will die during the movies are shown in white. Left schemes show the expected deformations of the active Ras cells (purple) and the neighbouring WT cells. Bottom right inset shows E-cad::GFP in Ras clones with strong apical constriction induced by activation of MyoII. Scale bars=10μm. **D:** Quantification of the probability of cell death for a given distance to clone boundaries (in cell rows). **** =p<10^-4^, ***=p<10^-3^, **=p<10^-2^, **-** =p>0.05, Fisher exact test with UAS-ras^V12^ (green bars). **E:** Evolution of the area covered by WT cells (one circle=one region surrounded by at least two clones). Bars are averages. p-value, t-test with UAS-ras^V12^ (green). **F:** Average *ras^V12^* cell apical area for the three first layers of cells near clone boundaries at the onset of the movie and after 700min. p-value, Mann-Whitney tests between the two time points, ****=p<10^-4^, cells are taken respectively from 4, 4 and 3 nota. Note that upon activation of MyoII, cells are already smaller at the onset of the movie as transcription was activated 8h before the onset of the movie (see Methods).

Next, we tested whether ERK downregulation was indeed necessary for neighbouring cell elimination. Overexpression of a secreted form of Spitz (*Drosophila* EGF) by *Ras^V12^* clones was sufficient to prevent neighbouring cell elimination (**Fig. 6C,D, movie S11**). This significantly slowed down the shrinkage of the area covered by WT cells (**Fig. 6E**) and the expansion of the *Ras^V12^* cell area which are close to the clone boundaries (**Fig. 6F,** *Ras^V12^* cell apical area near clone boundaries increased by 79% on average over 700min in controls, while only 24% for *Ras^V12^* and *sSpitz* expressing cells), which are the cells that should compensate for most of the area left by the WT dying cells[37]. This suggested that cell elimination through ERK downregulation contributed to pretumoural clone expansion and global replacement of the WT cells.

Finally, we wanted to make sure that WT cell elimination was indeed driven by mechanical cues rather than diffusive factors from the clones. Increasing contractility in *Ras^V12^* cells through the overexpression of an active form of MyoII (*UAS-sqhE20E21*[38]) was sufficient to prevent *Ras^V12^* cell expansion (**Fig. 6F,** average cell area decrease of 18%), and strongly downregulated WT cell elimination (**Fig. 6C-E**, **movie S11**), suggesting that *Ras^V12^* cell expansion is indeed necessary for WT cell elimination. Strong activation of the EGFR/ERK pathway can also upregulate the transcription of Argos, a secreted protein which feedbacks negatively on EGFR signaling through Spitz sequestration [39]. Therefore, elimination of WT cells could also be driven by an Argos-dependent ERK downregulation. However while downregulation of Argos in *Ras^V12^* clones by dsRNA overexpression completely prevented Argos accumulation in the clones, it did not abolish WT cell elimination near active Ras clones (**Fig. S4B,C, movie S11**). Those experiments confirmed that cell elimination was driven by mechanical cues rather than purely diffusive factors from the Ras clones.

Altogether we concluded that tissue compaction and/or a reduction of cell tension could rapidly downregulate ERK activity *in vivo*, which could trigger cell elimination through the upregulation of Hid. This modulation of ERK was responsible for cell elimination near *Ras^V12^* clones, which contributed significantly to clone expansion and could promote tissue invasion.

## Discussion

EGFR/ERK/Hid pathway has been previously involved in tissue homeostasis and cell number regulation through cell survival regulation [40, 41]. For instance, modulation of segment size in *Drosophila* embryo can also be adjusted by EGFR/Hid dependent death which is regulated by the limited source of Spitz/EGF ([41] and JP Vincent personal communication). Similarly, secretion of Spitz/EGF by a subset of cells and local activation of EGFR regulates the number of interommatidial cells in the fly retina by suppressing Hid activity both at larval stage[42-44] and during pupal development[18, 44, 45]. EGFR/Hid also regulates the number of glia cells in the embryo through limited amount of Spitz secreted by neurons[46]. All those studies focused on the modulation of EGFR/ERK by limitant extracellular ligands. Here, we show that ERK activity can also be modified by tissue mechanics *in vivo* which can modify cell survival. Modulation of ERK activity by mechanical stress and/or tissue density has been previously described in cell culture [30, 32, 33, 35, 47] and *in vivo* [34], however the functions of those mechanical modulations were not explored and most of those approaches remained correlative. Here, we provide the first evidence of a mechanical modulation of ERK playing an instructive role for cell survival/death during competitive interactions between two cell types *in vivo*. So far, most of the studies of mechanotransduction *in vivo* focused on the regulation of Hippo/Yap-Taz pathway [11] whose transcriptional outputs should act on hours timescale. Our results suggest that ERK modulation could act on cell survival in a few tens of minute (see **Fig.4**) either through the modulation of Hid activity by phosphorylation [17] and/or through transcriptional regulation [18]. While we found that tissue stress/compaction can modulate ERK activity and that part of the ERK dynamics correlated with tissue deformations, the complex spatiotemporal pattern of ERK activity in the pupal notum is very likely to be controlled as well by currently unknown patterning genes. EGFR/ERK modulation by mechanical stress may be required then to fine tune its activity and to coordinate in time and space cell elimination and to regulate the number of cells that will be eliminated. In that framework, a high rate of cell elimination would lead to higher cell spacing and/or an increase of tissue tension, which would feedback negatively on cell elimination through ERK activation. Further exploration of the mechanical modulations of ERK occurring *in vivo* in different tissues will be required to assess its contribution to tissue homeostasis, size regulation and morphogenesis. Interestingly, ERK was previously shown to modulate cell tension and tissue mechanics[29-31]. The mutual regulation of ERK and cell mechanics could be at the basis of some complex temporal dynamics and self-organizing properties of epithelial tissues *in vivo*, as previously observed in cell culture during collective cell migration[30].

While we still do not know which molecular effectors of ERK pathway are sensitive to mechanical stress, epistasis experiments suggest that modulation occurs upstream and/or at the levels of EGFR (see Fig. **S5 A-C**, **movie S12**). Accordingly, EGFR is mostly located on the apical side of the cells (**Fig. S5D**) which could be compatible with a modulation of EGFR concentration/activity by apical cell geometry and/or apical mechanical stress. Moreover, the correlation between tissue stretching/compaction and ERK activity could be explained by different parameters, including an effect of membrane tension, a direct readout of cell apical surface, and/or changes in cell volume affecting EGFR activation and/or ligand binding. The transient ERK downregulation observed after tissue severing and the correlation between compaction rate and ERK activity would suggest that ERK is sensitive to variations of cell geometry in time rather than absolute cell size/tissue density. Similarly, compressive forces rather than absolute tissue density were shown to be responsible for spontaneous MDCK cell elimination[6]. Further exploration of the single cell parameters correlating with ERK fluctuations will help to identify the relevant factors involved in compaction driven ERK downregulation and to elucidate the molecular mechanism of ERK mechanotransduction.

Finally, the contribution of compaction induced ERK inhibition to cell elimination and *Ras^V12^* clone expansion may be relevant for other competition scenarios and in pathological conditions. Accordingly, Yki activation in clones was also shown to trigger cell deformation and cell elimination in the pupal notum[37]. Elimination of cells mutant for the apico-basal polarity protein Scribble are also eliminated through a downregulation of EGFR/ERK[48], however this process is driven by a contact dependent ERK downregulation through the ligand Sas and the tyrosine phosphatase PTP10D. Finally, compaction driven ERK downregulation may also be relevant for the elimination of miss-pecified cells in *Drosophila* wing imaginal disc, which has been associated with an increase of contractility at the clone boundary leading to cell compaction within the clone[49]. Constitutively active mutant forms of Ras are present in one third of human cancers [50]. Our study suggests that downregulation of mechanical induced cell elimination could significantly slowdown the expansion of tumoural cells. The opposite effects of ERK on tumours (promotion of tumoural cell growth and survival, regulation of neighbouring cell resistance to mechanical stress) may explain the limited success of Ras/Raf/ERK targeted cancer therapies [50].

## Aknowledgements

We would like to thank members of RL lab and F Schweisguth for critical reading of the manuscript. We are also very grateful to F. Janody, M. Miura, C. Bökel, H.D. Ryoo, G. Jiménez, the Bloomington Drosophila Stock Center, the Drosophila Genetic Ressource Center, the Vienna Drosophila Ressource Center, and the Developmental Studies Hybridoma Bank for sharing stocks and reagents. We are also grateful to B Aigouy for the Packing Analyser software and members of J. Ellenberg group for MyPic macros for autofocus. LV is supported by a Post-doctoral grant “Aide au Retour en France” from the FRM (Fondation pour la Recherche Médicale, ARF20170938651), work in RL lab is supported by the Institut Pasteur (G5 starting package) and the ERC starting grant CoSpaDD (Competition for Space in Development and Disease, grant number 758457). Work in EM lab is supported by the European Research Council, The Swiss National Foundation and the Champalimaud Foundation.

### Authors contribution

EM and RL initiated the project. LV performed the wounding experiments, the cross-correlation analysis, the miniCic/caspase correlation and the local projection algorithm. FL performed the dpERK western blot. RL wrote the manuscript and performed all the other experiments and analysis. All the authors have commented and edited the manuscript.

### Competing interest

The authors declare no competing interest

## Star methods

Reagents and further details about protocols and macro can be shared upon request to Romain Levayer (http://romain.levayer@pasteur.fr).

### Fly stocks and clone induction

We list here the fly lines used for each figure as well as the time of heat shock when relevant and the days after clone induction (ACI).

| Figure | Genotype | Origin and RRID | Heat shock 37°C | Flybase ID |
| --- | --- | --- | --- | --- |
| 1b,<br>2b,S2a | *pnr-gal4 (III)* | Bloomington<br>(BDSC_3039) | - | FBst0003039 |
| 1b | *UAS-diap1 (III)* | Bloomington<br>(BDSC_6657) | - | FBst0006657 |
| 1b,c, 2b | *UAS-hid ds RNA (III)* | VDRC GD 8269 | 12 min, 3 days ACI | FBst0470995 |
| 1b,c | *UAS-rpr dsRNA (II)* | VDRC KK 101234 | 12 min, 3 days ACI | FBst0473107 |
| Not<br>shown | *UAS-grim dsRNA (II)* | VDRC GD 21830 | 12 min, 3 days ACI | FBst0454257 |
| 1 c,d | *hs-flp22;ubi-ECad::GFP,UAS-*<br>*mRFP; act<y+<Gal4* | L. Legoff (Marseille) | 12 min, 3 days ACI |  |
| 1f, 6a-f | *UAS-ras1<sup>V12</sup> (III)* | From [51] | 30 min, 1 week 18°C |  |
| 2a | *cic-Cic::GFP (Bac clone) (II)* | Bloomington<br>(BDSC_42267) |  | FBst0042267 |
| 2b,c,d<br>3c, S5a-c | *UAS-egfr dsRNA (II)* | VDRC KK 107130 | 12 min, 3 days ACI | FBst0478953 |
| Not<br>shown | *UAS-egfr dsRNA*<br>*TRiPJF01084 (III)* | Bloomington<br>(BDSC_31526) | 12 min, 3 days ACI | FBst0031526 |
| 2c,d<br>3c | *UAS-hEC::EGFR (II)* | Bloomington<br>(BDSC_58415) | 12 min, 3 days ACI | FBst0058415 |
| S1b,c | *tub-GFPBantamsensor/Cyo (II)* | From [16] |  | FBtp0017244 |
| S1c,d,e | *UAS-dsRed Bantam sponge*<br>*(III)* | From [52] | 12 min, 3 days ACI | FBtp0073033 |
| S1b | *ubi-Ecad::RFP (II)* | From [53] |  | FBal0295303 |
| S2a | *UAS-pvr dsRNA (II)* | VDRC KK 105353 |  | FBst0477180 |
| S2a | *UAS-fak dsRNA (III)* | VDRC GD 108608 |  | FBst0480418 |
| S2a | *UAS-src42A dsRNA (II)* | VDRC KK100708 |  | FBst0472581 |
| S2a | *UAS-src64B dsRNA (I)* | VDRC GD 12263 |  | FBst0461056 |
| S2b | *UAS-pi3kCAAX (I)* | Bloomington<br>(BDSC_8294) | 12 min, 3 days ACI | FBst0008294 |
| S2c | *UAS-raf<sup>GOF</sup> (III)* | Bloomington<br>(BDSC_2033) | 12 min, 2 days ACI | FBst0002033 |
| 3b-f, S3,<br>4a-c, 5a-<br>j, 6a,b | *tub-miniCic::mCherry (II)* | This study |  |  |
| 3c | *hs-flp22 ; endo-Ecad::GFP ,*<br>*miniCic::mCherry ;*<br>*act<cd2<G4, UAS-*<br>*eGFP::mCd8* | Bloomington<br>(BDSC_60584) and<br>this study | 12 min, 2 days ACI | FBst0060584 |
| 3c,f<br>5a-k | *endo-Ecad::GFP,tub-*<br>*miniCic::mCherry* | Bloomington<br>(BDSC_60584) and<br>this study |  |  |
| 4 a-c | *UAS-scat3::myc (II)* | From [28] |  | FBtp0021422 |
| 1f, 6a-e | *hs-flp22; endo-Ecad::GFP, tub-*<br>*gal80ts ; act<y+<gal4 UAS-*<br>*nlsRFP/TM6b* | From [54] | 30 min, 1 week 18°C |  |
| 6c-f | *UAS-sSpitzCS , UAS-rasV12*<br>*(III)* | Bloomington<br>(BDSC_63134)<br>recombined with<br>*UAS-ras<sup>V12</sup>* | 30 min, 1 week 18°C | FBst0063134 |
| 6c-f | *UAS-sqhE20E21 (II) ; UAS-*<br>*rasV12 (III)* | From [55] combined<br>with *UAS-ras<sup>V12</sup>* | 30 min, 1 week 18°C | FBal0246103 |
| S4a,b | *UAS-argos dsRNA (II) ; UAS-*<br>*rasV12 (III)* | VDRC GD 47181<br>combined with *UAS-*<br>*ras<sup>V12</sup>* | 30 min, 1 week 18°C | FBst0467120 |
| S4d | *egfr-EGFR::GFP (MiMIC) (II)* | Bloomington<br>(BDSC_61765) |  | FBst0061765 |

### Immunostaining

Dissection and immunostainings of nota were performed as indicated in [56] with standard formaldehyde fixation and permeabilisation/washes in PBT 0.4% Triton. Immunostaining of wing imaginal discs and eye imaginal discs were performed on L3 wandering stage larvae and fixed in 4% formaldehyde and permeabilised/washed in PBT. Immunostaining of embryos were performed using standard formaldehyde and heptane/methanol protocol, with permeabilisation in PBS Tween (0.05%). The following antibodies/markers were used: rat anti E-cad (DCAD2 concentrate, DSHB, 1/50), guinea pig anti Hid (1/50, strepatavidin amplified, gift of Don Hyong Ryu), mouse anti EcR (DSHB Ag10.2 concentrated, 1/100), rabbit anti dpERK (Cell signaling, #4370, 1/100), chicken anti GFP (abCam, #13970, 1/500), chicken anti mCherry (abCam, #205702, 1/200), rabbit anti-cleaved Dcp-1 (Cell Signaling, #9578, 1/50), mouse anti-Argos (DSHB concentrated, 1/50), rabbit anti EGFR (SantaCruz, #33741, 1/50). Secondary antibodies were all Invitrogen secondary antibodies produced in goat with Alexa 488,555 or 633. Dissected nota were mounted in Vectashield with DAPI (Vectorlab) and stained embryos were mounted in Aquapolymount media. They were imaged on a Leica confocal SP8 or a Zeiss lsm880 using a 63X oil immersion objective N.A. 1.3. Images shown are maximal z-projection containing adherens junction plane.

### Design of the miniCic sensor

The C-ter part of Capicua containing the C2 and C1 domains but lacking the DNA binding domain (HMG box) was PCR amplified from a cDNA clone (DGRC clone FI04109, starting at base number 2687, sequence: *TCCGCCTCCGGAGGGGGCGTGGTC*) adding a NLS site in N-ter right after the ATG. The PCR product was fused with mCherry through fusion PCR. The fusion product was inserted in pC4-tub-gal80 using the NotI and XbaI sites after removal of the Gal80 through digestion, purification and ligation. The construct was validated by sequencing and injected by Bestgene (P-element insertion). Further details about the construct can be sent upon request.

### Western blot

S2R+ cells were cultured in Schneider’s *Drosophila* Medium with 10% fetal bovine serum, penicillin and streptomycin. Cells were plated at a density of 2x10^6^ cells into twelve well plates and incubated at 25°C. 24h after plating, they were treated with Trametinib 10μM dissolved in DMSO or with DMSO alone (1/1000). Cellular extracts were prepared at indicated time post treatment and analysed by immunoblotting. The following primary antibodies were used: phospho-Erk(Thr202/Tyr204) (1/1000, Cell signaling, #4370), total-Erk (1/1000, Cell signaling, #4695) and alpha-tubulin(1/5000, DSHB, 12G10 concentrated).

### Analysis of adult thorax defects

Adult thorax defects were analysed on adult female thorax raised at 25°C. For **Fig.1B** we measured the width of the midline divided by the averaged width of the two adjacent SOP rows along a line connecting the two aDC macrochaetaes (log2 scale). For **Fig. 2B** we measured the width of the *pnr* domain (distance between the two aDC macrochaetaes) divided by the total width of the thorax (distance between the two pSA macrochaetae) and plotted the log2 of the ratio.

### Imaging of haemocytes and drug treatment.

Primary culture of haemocytes was performed by bleeding ten larvae expressing *tub-miniCic::mCherry* that were first washed in ethanol, dried on a piece of paper and resuspended in 200μL of S2 medium on a square of parafilm. After bleeding with forceps each larvae, the 200μL were deposed in a MaTek petri dish (P35G-1.5-10-C) and we let the cells sediment and adhere for 30min. The media was then supplemented by 2mL of S2 medium. The dish was placed on a LSM800 point scanning confocal (63X oil N.A. 1.3) and acquisition was launched with a transmitted light channel and mCherry channel (one z-stack/min). Trametinib (Santa Cruz, 10mM stock in DMSO, final concentration 10μM) or DSMO (1/1000) were added in the medium at the onset of the movie. Nuclear miniCic signal was measured by tracking manually the nucleus using the transmission light channel on Fiji, measuring mCherry intensity and normalising intensity by intensity at t0.

### Notum live imaging, image processing and cell death analysis

Notum live imaging was performed as described previously[9]. Pupae were collected 48 or 72h after clone induction and dissected 16-18h after pupae formation (APF). Pupae were dissected and imaged on a confocal spinning disc microscope (Till photonics) with a 40X oil objective (N.A. 1.35) using tile imaging (6 to 12 tiled positions) or a point scanning confocal microscope Leica SP8 with a 63X objective (N.A. 1.3), or a Zeiss LSM800 or a LSM880 equipped with a fast Airyscan using an oil 40X objective (N.A. 1.3), Z-stacks (1 μm/slice), every 5min using autofocus at 25°C. The autofocus was performed using E-cad::GFP plane as a reference (Tillphotonics and Leica LAS provided option) or the autofluorescence of the cuticle in far red (using a Zen Macro developed by Jan Ellenberg laboratory, MyPic). Movies were performed in the nota close to the scutellum region containing the midline region and the aDC and pDC macrochaetae. Movies shown are maximum projections or adaptive local z-projection (see below for details). Total duration was always 700min (except for Fig. 6A, 600min). For imaging of *UAS-ras^V12^* clones, the cross and the progeny were kept at 18°C, and the pupae were switched to 29°C 8 hours prior to the movie for conditional activation (clones induced by the following lines: *hs-flp22; endo-Ecad::GFP, tub-gal80ts; act<y+<gal4 UAS-nlsRFP/TM6b*) and imaged at 29°C. The midline region covers all the cells surrounded by the two most central lines of sensory organ precursors (SOP, located at the end of the movies). Every cell extrusion event was localised manually and marked using Fiji. The probability of extrusion was obtained by dividing the total number of extruding cells in the clones divided by the initial number of cells in the clones or by doing the same for each cell layer next to Ras clones. Values from the control clones (ayGal4 alone) are coming from [9] as part of the clonal experiments were performed during the same period. In the figures, cells were marked as dying cells if they died or if at least one of their daughter cells died before the end of the movie.

For the quantification of the area covered by WT cells during competition with Ras, regions of WT cells surrounded by at least two Ras clones were defined on Fiji and their area was measured at the onset of the movie and after 700min. Ras cell area was obtained after segmentation of the images at the onset of the movie and after 700min using packing analyser Fiji plugin[57]. We measured apical area of the three first layer of cells near clone boundary.

### Adaptive local z-projection and analysis of miniCic::mCherry

All the image of miniCic::mCherry in the notum were obtained through a custom made adaptive local z-projection procedure wrote on Matlab (to avoid projection of the autofluorescence signal of the cuticle and projection of more basal plane). For every time point, a reference channel was used (here E:cad::GFP), the image was subdivided in 40*40px (1px=0.18μm) square and the plane with the highest standard deviation was selected. The search window was moved in x and y 10px by 10px. The obtained z-profile was then smoothened in x and y using a Gaussian blur (2px width). miniCic signal was then obtained by calculating the median of the intensity over seven planes centered on a reference plane located 6μm basally to the local E-cad reference plane. Note that similar results could be obtained by using maximum projection instead of the median. For every movie, we checked on the raw files that no nuclear signal of miniCic was lost because of projection errors.

### Nuclear miniCic quantification and Scat3 imaging

Live pupal nota expressing UAS-Scat3 with *pnr-gal4* driver and miniCic were imaged on a LSM880 point scanning confocal (one stack every 5min or 10min). Images were pretreated with local z-projection (see above, YFP used as a reference channel, average of 7 planes around the plane of reference). Image ratios of YFP over CFP fluorescent channel were then calculated (FRET signal). Cells showing a clear caspase activation (low YFP/CFP ratio) were selected manually. The miniCic channel was also cut according to those positions and smaller stacks of 15 time frames were created (1 stack/10 min). Positions of the nuclei were selected manually over the 15 frames for all the selected cells and both YFP/CFP ratio and miniCic mean intensity values were quantified in boxes of 2μm squares centered on those positions. The average profiles were obtained by aligning the curves on the point of maximum inflexion of the FRET signal (local minimum of the 2^nd^ derivative). Each single curve was then normalised by the values at the 3 first time points (70 to 50 min before caspase activation). For miniCic, the intensity value at t0 was also subtracted. All the curves were then averaged. Non-ambiguous ERK downregulation (as mentioned in the text) corresponds to curves with at least a 5% increase of miniCic signal over one hour prior to caspase activation. The same procedure was performed for the control curve on randomly selected cells that do not activate caspase in same region and time developmental window of the caspase activating cells. For those curves, alignment was made at T0 (onset of the tracking).

### Particle Image Velocimetry, cross-correlation and tissue wounding

For **Fig. 5**, we used a PIV algorithm based on a custom made Matlab routine as described in[58]. Parameters for this PIV were analysis boxes of 64 pixels (11.5x11.5 μm ∼4-5 cells), 50% overlap boxes. PIV displacements at time t were determined based on images t-5 and t+5min. Displacement maps were spatially filtered to remove noise coming from outliers.

Quantification of miniCic mCherry signal (**Fig. 5A-G**) are median values of mCherry signal over the analysis boxes of the PIV after performing local projection. For each notum, the region of interest (zones of maximum deformation) were subdivided in 25 squares of analysis (50% overlap). Compaction rate maps were calculated as minus the “divergence” function of Matlab on the PIV vector fields. miniCic signals were smoothen over ten time frames before calculating the derivative, compaction rate were smoothen the same way. The average value was subtracted from each function to obtain variations around 0 value. Maps of cross-correlation and normalised cross-correlation between compaction rate and derivative of miniCic levels were then calculated with the “xcorr” Matlab function either without or with the ‘coef’ option (non-normalised and normalised cross-correlation). Quantification of cross-correlations were established in stretching areas for time periods of 2 hours. Number of nota rather than number of square of analysis was used to calculate the s.e.m‥ To obtain control cross-correlation curves, we computed the normalised cross-correlation of miniCic and compaction rates in regions with no obvious deformations. We also checked that there were no correlation between compaction rate of the stretched area and miniCic signals from regions outside the stretch area to make sure that the miniCic variations were specific of the zones of deformation (data not shown). Single cell tracking and segmentation (**Fig. 5H,I**) was performed using packing analyser Fiji plugin[57] and by manually tracking the corresponding miniCic nuclei for each cell.

Notum wounding was performed as described previously[9], using exposure at full power with an Argon laser (488 and 458nm) and a 405nm diode in the wounded region over 3min. For experiment in **Fig. 5J,K**, tissue square dissection was performed using a pulsed 355nm laser (Teem photonics, 20kHz, peak power 0.7kW) controlled by the ILas pulse system (Gataca systems) on an inverted Nikon eclipse Ti2 microscope (plan fluor 40X 1.3 N.A, oil) equipped with a Yokogawa CSU-W1 spinning disc system and a prime95 sCMOS camera (Photometrics). The region of ablation was a 400x400px square (110x110μm) with a band of 10px width and 10 repetitions (AOTF 35%). Relaxation was imaged using z-acquisitions in green and red channels every 2min combined with an autofocus between each time point. Imaged shown are local z-projections. Quantification of cell area was performed in Tissue analyser[57], intensity of miniCic was quantified after tissue drift correction (Fiji plugin StackReg), bleaching correction (on the full movie) and removal of the background (rolling ball radius 50px).

For **Fig. 6,** PIV analysis was performed as described previously[9], using MatPIV toolbox on Matlab with 64px windows with 50% overlap and a rolling window of time averaging of 9 frames. Compaction rate is defined as “*- Divergence”* of the vector field (calculated on Matlab). Averaged compaction rates were calculated by averaging on one PIV window the compaction rate over the full movie (700 min). The same squares were used to quantify miniCic signal at the last time point of the movie excluding any zone that partially overlapped with the Ras clones. The mean miniCic signal was then calculated for each bin of compaction rate values and the full data set was used to calculate a Pearson correlation coefficient.

### Statistics

Unless specified, data were not analysed blindly. No specific method was used to predetermine the number of samples. Error bars are standard error of the mean (s.e.m.). p-value are calculated through t-test if the data passed normality test (Shapiro-Wilk test), or Mann-Whitney test if the distribution were not normal. For proportion (death probability), the error bars are 95% confidence interval and p-value are calculated through a Fisher exact test. Statistical tests were performed on Graphpad.

---

**Figure S1:**
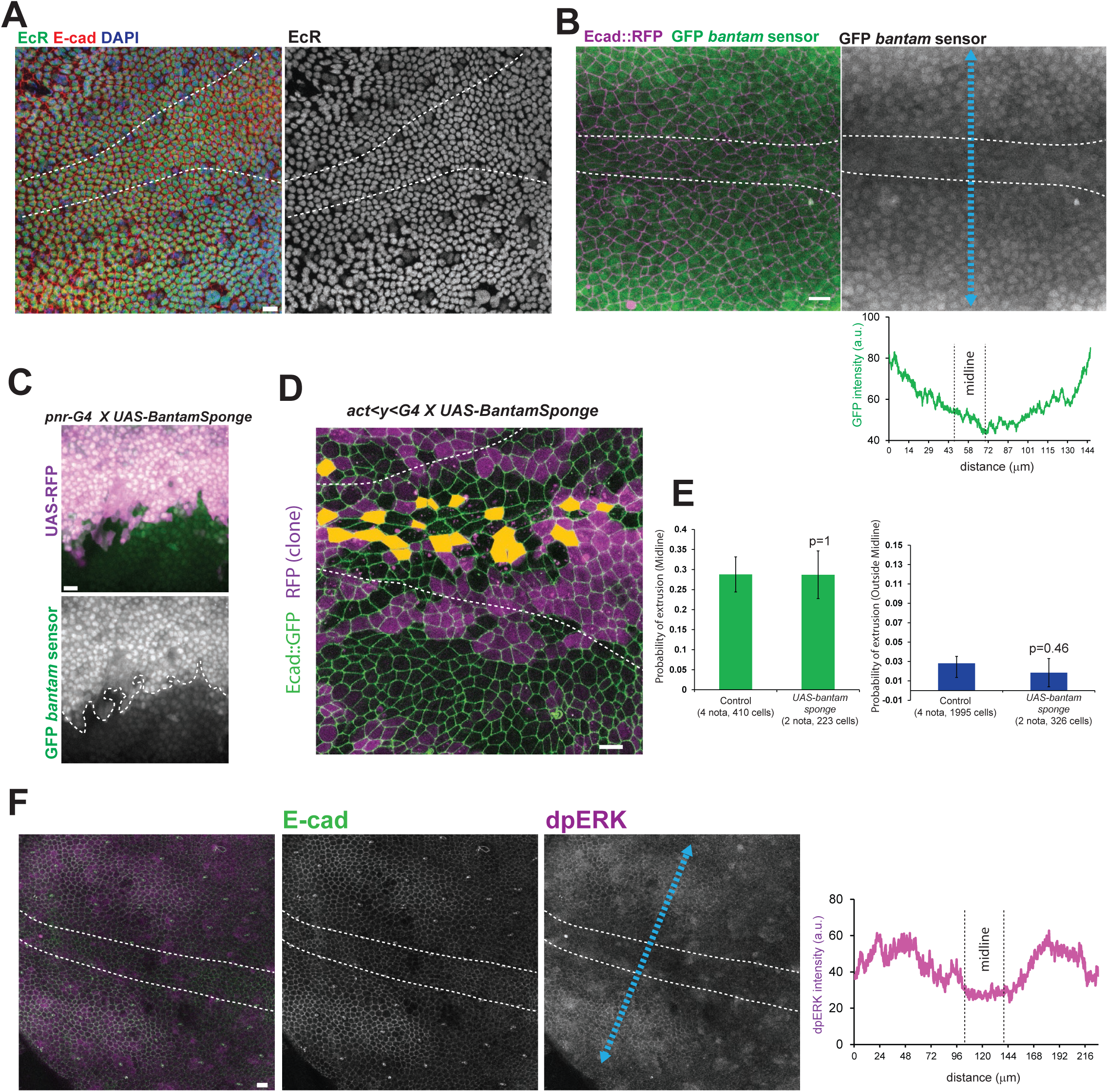
Bantam and Ecdysone are not responsible for cell death in the midline (associated with movie S2) **A:** z-projection of an immunostained pupal notum for Ecdysone receptor (EcR, green), DAPI (blue) and E-cad (red). The midline is encompassed by the dashed lines. Scale bar=10μm. **B:** z-projection of a live pupal notum expressing ubi-Ecad::RFP (magenta) and a GFP *bantam* sensor (green, low GFP= high *bantam*), midline region marked by white dashed lines. Intensity profile of the GFP along the blue line is shown below. Scale bar=10μm. **C:** z-projection from a live pupal notum expressing UAS-BantamSponge and a RFP in the *pnr* domain (magenta) and the GFP bantam sensor (bottom). GFP accumulates in the *pnr* domain compared to more lateral domains, suggesting that Bantam activity is indeed reduced by the sponge. Scale bar=10μm. **D:** z-projection of live a pupal notum expressing *ubi-Ecad::GFP* (green) with Gal4 expressing clones (RFP, magenta) expressing a *bantam* sponge. The midline is delineated by white dashed lines. Orange cells are the clonal cells that will die over the course of the movie (700 min). See **Fig. 1C** for control. Scale bar=10μm. **E:** Quantification of the probability of cell elimination in the clones in the midline region (left) and outside the midline region (right). pvalues are Fisher exact tests with the control condition (same as **Fig. 1C,D**). **F:** z-projection of an immunostained pupal notum for E-cad (green) and dpERK (magenta). The midline is encompassed by the dashed lines. Scale bar=10μm. Intensity profile of dpERK along the blue line is shown on the right.

**Figure S2:**
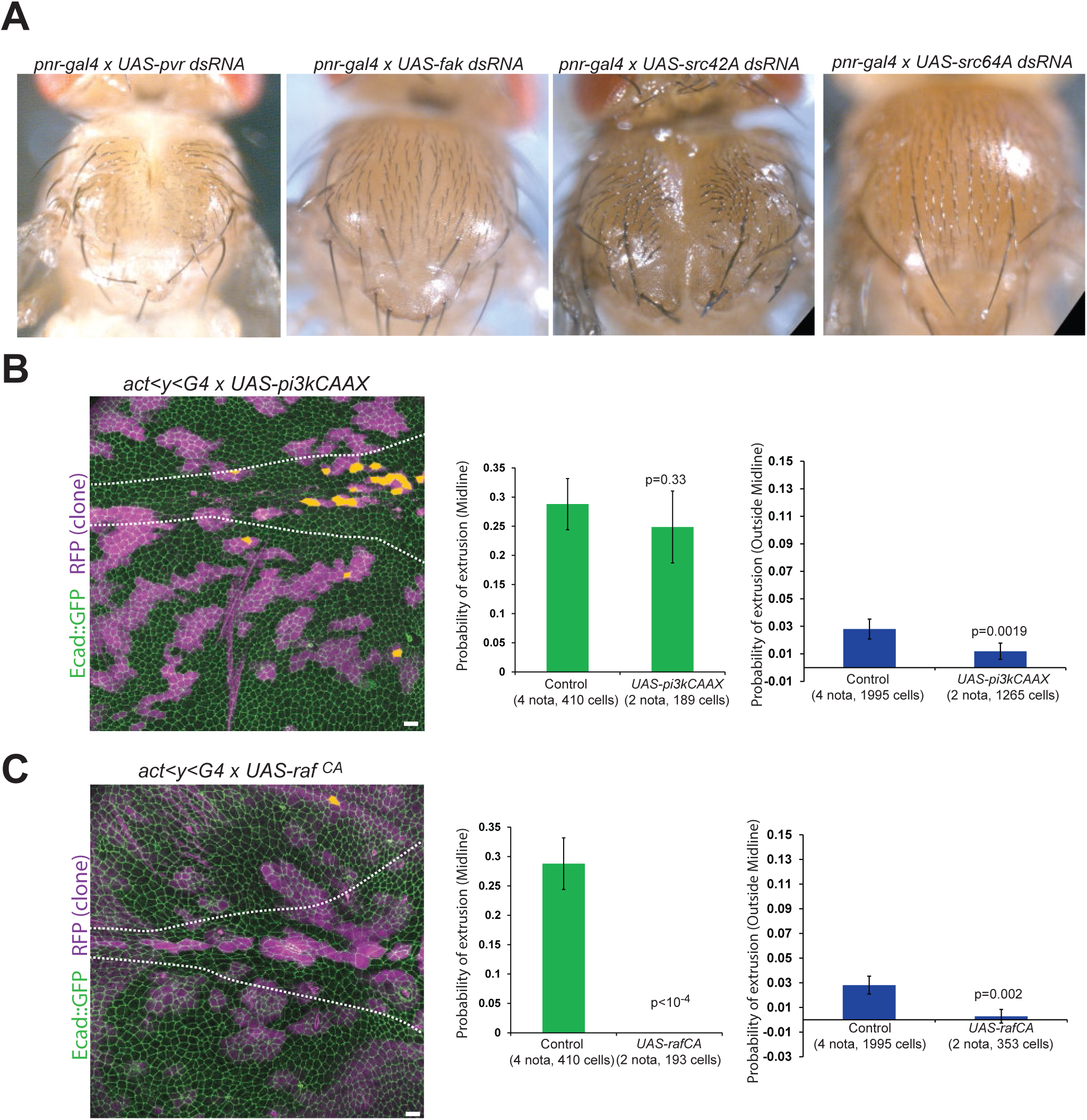
Cell survival is regulated by Raf, not PI3K, PVR, FAK, Src42A or Src64B (associated with movie S2) **A:** Representative adult thoraxes upon PVR, FAK, Src42A or Src64B downregulation by RNAi using the *pnr-gal4* driver. Compare with **Fig. 1B** (*UAS-lacZ*) for control (minimum of 10 thorax per condition). **B,C** z-projection of live pupal nota expressing *ubi-Ecad::GFP* (green) with Gal4 expressing clones (RFP, magenta) expressing active PI3K (*UAS-pi3kCAAX*, top) or active Raf (*UAS-raf^CA^*, bottom). The midline is delineated by white dashed lines. Orange cells are the clonal cells that will die over the course of the movie (700 min). See **Fig. 1C** for control. Scale bars=10μm. Right: Quantification of the probability of cell elimination in the clones in the midline region (left, green) and outside the midline region (right, blue). p-values are Fisher exact tests with the control condition (same as **Fig. 1C,D**). Note that we previously showed the efficiency of PI3K pathway activation upon *pi3kCAAX* expression exactly in the same condition previously[54].

**Figure S3:**
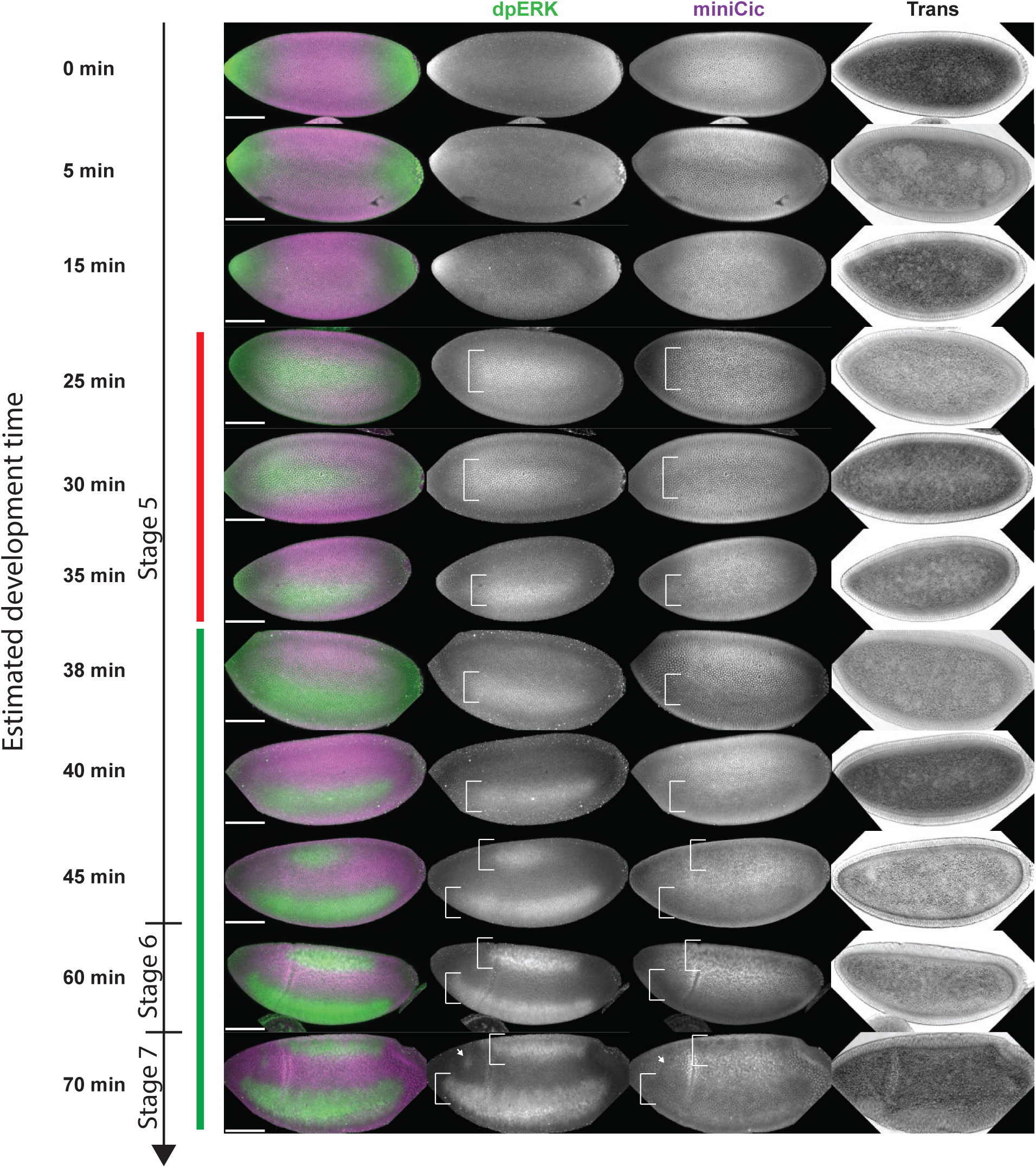
miniCic accurately reports levels of ERK activation in the embryo. z-projections of immunostained embryos at the cellularising (stage 5) and early gastrulating stage (stage 6,7). dpERK in green, anti mCherry (for miniCic::mCherry) in magenta. Transmitted light on the right. Development time (after onset of cellularisation) is estimated by the size of the invaginating membranes compared to movies of cellularising embryos (not shown). White brackets show the ventral lateral zone of activation of ERK and the dorsal zone of activation. The red bar shows the first stages of dpERK activation where there is no obvious local downregulation of miniCic. At later stages, the dpERK pattern is well reflected by the absence of nuclear miniCic (brackets and white arrows). Scale bars=100μm.

**Figure S4:**
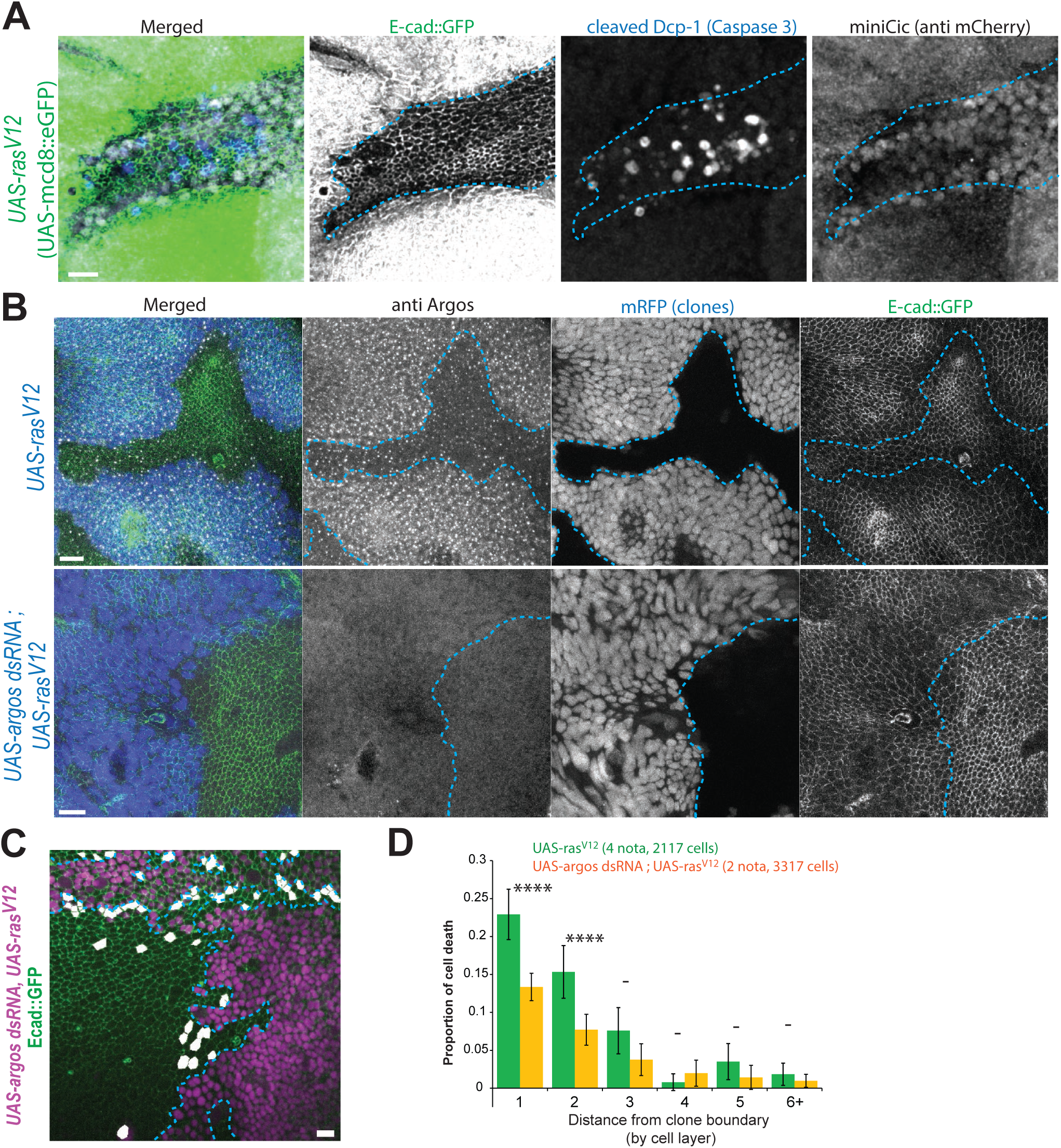
Cell elimination near Ras activated clones is not purely Argos dependent (associated with movie S11) **A:** z-projection in the notum near activated Ras clones (Green, mCD8::eGFP) showing apoptotic WT cells (blue, cleaved Dcp-1) in the region of low ERK activity (accumulation of nuclear miniCic). Blue doted lines show clone contours. Scale bar=10μm. **B:** z-projection in E-cad::GFP (green) nota near clones overexpressing active Ras (*UAS-ras^V12^*, blue=UAS-mRFP, top) or active Ras and a dsRNA targeting Argos (*UAS-argos dsRNA; UAS-ras^V12^*, blue=UAS-mRFP, bottom) stained for Argos (grey) 24h after activation at 29°C. Blue doted lines show clone contours. Argos accumulation in active clones is lost upon dsRNA expression. Scale bars=10μm, representative of 3 nota for each. **C:** z-projection of a live pupal notum expressing endo-Ecad::GFP (green) upon conditional induction of *Ras^V12^* and *argos-dsRNA* in clones (RFP, strong magenta signal, blue dashed lines=clone contours). The cells that will die during the movie are shown in white. Scale bar=10μm. **D:** Quantification of the probability of cell death for a given distance to clone boundaries (cell rows). Control (green bars) come from Fig. 6C,D. **** =p<10^-4^, **-** =p>0.05, Fisher exact test with *UAS-ras^V12^* (green bars). Although there is a diminution of the rate of elimination, there is still a strong increase of cell death near the clones (contrary to *UAS-sSpitz* or *UAS-sqhE20E21*, Fig. 6D).

**Figure S5:**
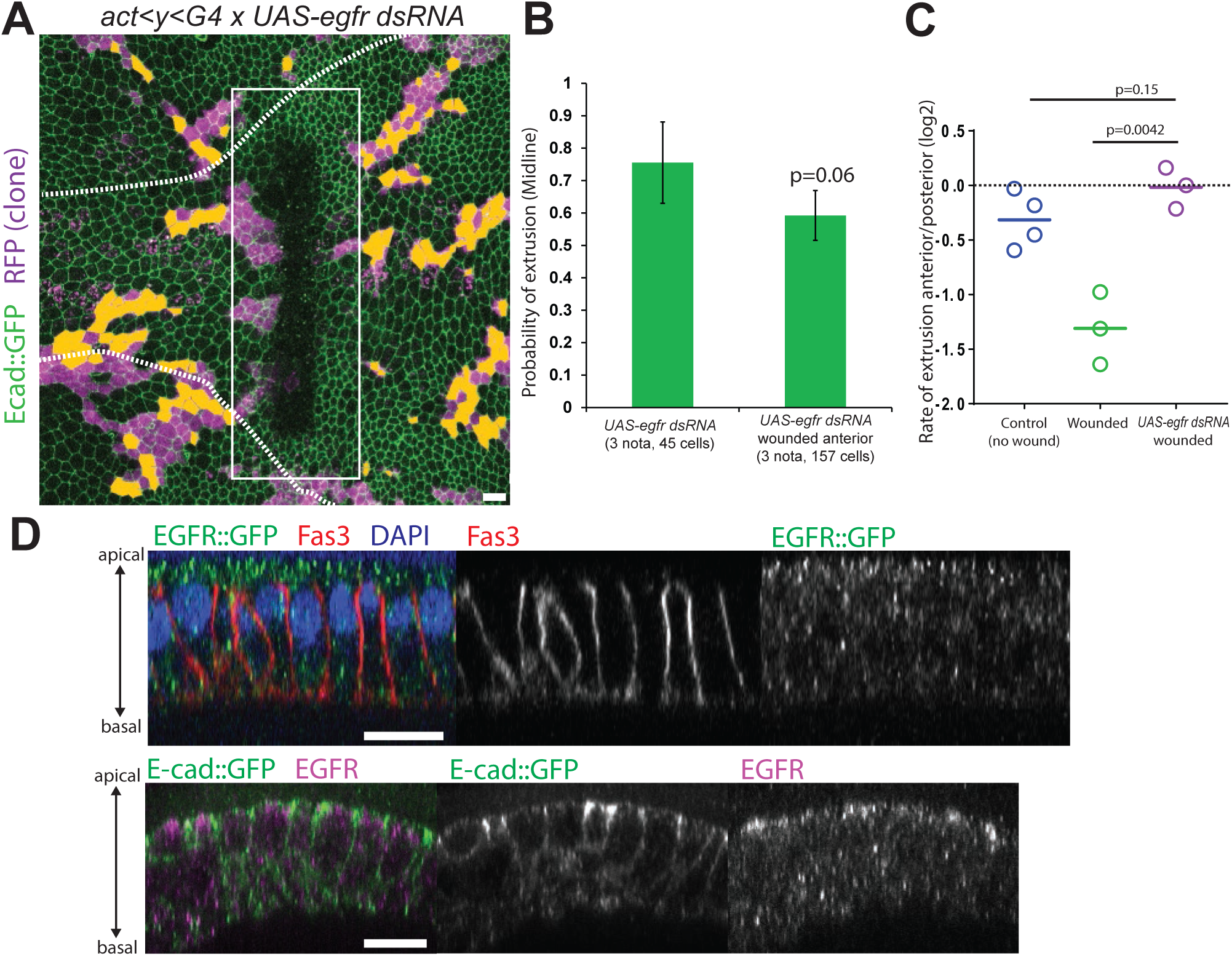
Mechanical sensitivity of ERK pathway is upstream and/or at the level of EGFR (associated with movie S12) **A:** z-projection of a live pupal notum expressing *ubi-Ecad::GFP* (green) with Gal4 expressing clones (RFP, magenta) where EGFR is downregulated (*UAS-egfr dsRNA*) after laser wounding (white rectangle, anterior on the left side, posterior on the right). The midline is delineated by white dashed lines. Orange cells are the clonal cells that will die over the course of the movie (700 min). Scale bar=10μm. **B:** Quantification of the probability of cell elimination in the clones in the midline region anterior and posterior to the wound. p-value is Fisher exact test. **C:** Quantification of the relative rate of cell extrusion in the midline region anterior and posterior to the wound in controls (no wounding), upon wounding (middle), and in wounded nota after depletion of EGFR in clones. One circle=one notum. Note that the data for the Control and Wounded are coming from [9]. While wounding normally induced a reduction of cell elimination anterior to the wound relative to the posterior region (since cell stretching is induced anterior to the wound because of the global tissue flow), there was however no apparent difference in the rate of cell elimination anterior and posterior to the wound in EGFR depleted clones (see p-values, t-tests). This was suggesting that EGFR depleted clones were insensitive to stretching, suggesting that stretching acts at the level or upstream of EGFR. **D:** Lateral view of cells in the pupal notum stained for Fasciclin 3 (Fas3, red) and GFP (EGFR::GFP trap, green) (top) or (bottom) EGFR (magenta) and GFP (endo-Ecad::GFP, green). Scale bars=10μm.

## Supplementary movie legends

**Movie S1: Hid downregulation prevents cell extrusion.** Related to Figure 1.

z projections of 16–18 hr APF pupal nota expressing *ubi-Ecad::GFP* (green) and Gal4 UASRFP clones (magenta) in control (*UAS-lacZ*, left), upon Hid downregulation in clones (*UAS-hid dsRNA*, middle) and Rpr downregulation in clones (*UAS-rpr dsRNA*, right). The midline is encompassed by the white lines and the extruding cells in the clone are marked in yellow at the beginning of the movie. Anterior on the left, posterior on the right. Scale bars=10 μm

**Movie S2: Midline cell extrusion is regulated by Raf, not bantam nore PI3K.** Related to Figure S1 and S2.

z projections of 16–18 hr APF pupal nota expressing *ubi-Ecad::GFP* (green) and Gal4 UASRFP clones (magenta) overexpressing Bantam sponge (*UAS-bantam sponge*, left), an active from of PI3K (*UAS-pi3k^CA^*, middle) or an active form of Raf (*UAS-raf^CA^*, right). The midline is encompassed by the white lines and the extruding cells in the clone are marked in yellow at the beginning of the movie. Anterior on the left, posterior on the right. Scale bars=10 μm

**Movie S3: EGFR downregulation is necessary and sufficient for cell extrusion in the pupal notum.** Related to Figure 2.

z projections of 16–18 hr APF pupal nota expressing *ubi-Ecad::GFP* (green) and Gal4 UASRFP clones (magenta) upon EGFR downregulation in clones (*UAS-egfr dsRNA*, left) and upon overexpression of hsEGFR::GFP (UAS-EGFR::GFP, right, clones are marked by the strong membrane GFP signal, note that the movie was filtered to obtain similar intensity ranges for E-cad and EGFR::GFP in the clones). The midline is encompassed by the white lines and the extruding cells in the clone are marked in yellow at the beginning of the movie. Anterior on the left, posterior on the right. Scale bars=10 μm.

**Movie S4: miniCic dynamics in haemocytes upon MEK inhibition.** Related to Figure 3.

Primary culture of haemocytes expressing miniCic::mCherry upon treatment with 10μM Trametinib (MEK inhibitor, left) or DMSO (right) at the onset of the movie. Scale bar=10μm.

**Movie S5: miniCic dynamics in the pupal notum.** Related to Figure 3.

Local z-projections of a live pupal notum expressing endo-Ecad::GFP (green) and miniCic (magenta and right). Time is indicated in hours after pupal formation. Scale bar=10μm.

**Movie S6: ERK downregulation precedes caspase activation.** Related to Figure 4

Local projection of two nuclei of the midline in a living pupal notum expressing miniCic (left) and Scat3 (FRET signal, right, red/orange=low caspase activity, blue-purple= high caspase activity). Scale bar=10μm.

**Movie S7: Fluctuations of miniCic in a WT notum during a stretch event orthogonal to the AP axis.** Related to Figure 5.

Left, miniCic mcherry fluorescent signal (local projection), middle, inverted endoCad::GFP signal (local projection), right, overlay of compaction rate (blue=stretching, red=compaction) and PIV maps all performed on a 16-18h APF notum. Black large rectangle on the PIV panel indicates the 30 frames used to analyse correlation between miniCic derivative and compaction rate. Red rectangle underlines the zone of stretching used for the analysis. Anterior on the right, posterior on the left. Scale bar= 20 μm. Color map and arrows scales are the same as indicated in **Figure 5A**.

**Movie S8: miniCic fluctuations during ectopic stretching induced by laser tissue wounding.** Related to Figure 5.

Left, miniCic mcherry fluorescent signal (local projection), middle, inverted endoCad::GFP signal (local projection), right, overlay of compaction rate (blue=stretching, red=compaction) and PIV maps on a 16-18h APF notum. The wound was done 10 minutes before the first image of this movie (low intensity rectangle on the left). Black large rectangles on the PIV panel indicates the 30 frames used to analyse correlation between miniCic derivative and compaction rate. Red rectangle underlines the zone of stretching used for the analysis. Anterior on the right, posterior on the left. Scale bar= 20 μm. Color map and arrows scales are the same as indicated in **Figure 5C**.

**Movie S9: miniCic increases upon tissue square severing.** Related to Figure 5.

Local z-projections of a live pupal notum (30h APF) expressing endo-Ecad::GFP (green, middle) and miniCic (magenta, right), upon severing of a 400x400px square of tissue with a pulsed UV laser (blue rectangle, ablation between 6 and 8 minutes). miniCic accumulates transiently in the nuclei of the isolated tissue. Anterior on the right, posterior on the left. Scale bar=10μm.

**Movie S10: ERK is downregulated in compacted regions near *Ras ^V12^* clones.** Related to Figure 6.

Local z-projection of a pupal notum expressing endo-Ecad::GFP (green) and miniCic (magenta, right panel) upon induction of *Ras^V12^* in clones (UAS-RFP, strong magenta). Note that the blurry rapid fluctuating signals are wandering macrophages below the tissue. Anterior on the left, posterior on the right. Scale bar=10μm.

**Movie S11: Activation of ERK or stretching of the neighbouring cells prevents cell elimination and slow down *Ras ^V12^* clones expansion.** Related to Figure 6 and Figure S4.

Local z-projections of pupal nota expressing endo-Ecad::GFP (green) upon induction of *Ras^V12^* in clones (UAS-nlsRFP, magenta, top left panel), upon induction of *Ras^V12^* and secreted Spitz (active *Drosophila* EGF, magenta top right panel), upon induction of *Ras^V12^* and active MyoII (UAS-sqhE20E21, magenta bottom left panel), and upon induction of *Ras^V12^* and downregulation of Argos (UAS-argos dsRNA, magenta bottom right panel),. Dying cells are marked in white prior to their extrusion. Anterior on the left, posterior on the right. Scale bars=10μm.

**Movie S12: EGFR depleted cells are not rescued by stretching.** Related to Figure S5.

z projection of a pupal notum expressing *ubi-Ecad::GFP* (green) and Gal4 UAS-RFP clones (magenta) where EGFR is depleted (*UAS-egfr dsRNA*) after laser wounding (white rectangle). The midline is encompassed by the blue lines and the extruding cells in the clone are marked in yellow at the beginning of the movie. Anterior is on the left, posterior on the right. Scale bar=10 μm

